# Loss of *lmx1ba* drives premature osteoarthritis through disruption of skeletal homeostasis

**DOI:** 10.64898/2026.04.27.720994

**Authors:** Joanna J. Moss, Felix Bowers, Joan Chang, Anthony Devlin, Stephen Cross, Elis Newham, Emily J. Rayfield, Jon D. Lane, Chrissy L. Hammond

## Abstract

Osteoarthritis is increasingly recognised as a disease of failed integration across the whole joint unit; however, the mechanisms that co-ordinate tissue integrity from development through to adult homeostasis remain largely unresolved. The LIM-homeodomain transcription factor LMX1B is a key determinant of embryonic skeletal patterning, but how it functions to regulate skeletal integrity in the mature skeleton is unknown. Recently, LMX1B was identified as a key driver of osteoarthritis. Here we show that loss of *lmx1ba* in zebrafish causes premature and progressive severe osteoarthritic pathology in adult spines and jaws, despite largely normal early skeletal patterning, revealing a conserved and continuous requirement for *lmx1ba* in joint maintenance beyond development. At a cellular level, loss of *lmx1ba* decouples osteoblast and osteoclast-mediated remodelling leading to bone overgrowth, heterogeneity of bone properties causing increased incidence of spontaneous fractures, and progressive abnormalities in spine morphology. In parallel, we observe degeneration of the intervertebral disc and dysregulation of the proteome and glycosaminoglycans indicative of disrupted extracellular matrix and a breakdown of the coordinated regulation of hard and soft tissue interfaces, which at the organismal level leads to altered joint performance. Notably, degeneration is restricted to mobile joints, and is not observed in cranial sutures, demonstrating a selective requirement for *lmx1ba* in mechanically active tissues. These changes are consistent with a model of spatially disrupted matrix properties that, under cyclic loading, promote progressive tissue damage. Our findings support a model in which continued expression of *LMX1B* in adulthood is required to maintain joint structures throughout life.

## INTRODUCTION

LMX1B belongs to the LMX group of the LIM-homeodomain family, a diverse family of regulatory proteins which are characterised by the presence of a DNA binding homeodomain and two protein binding LIM domains (LIM-A and LIM-B)^1,2^. Through these domains, LMX1B is able to both directly and indirectly regulate gene expression across a wide range of tissues such as the skeleton, brain and kidneys, and plays a critical role in early vertebrate development^3^. While its function in limb patterning is well established, emerging evidence suggests that LMX1B may contribute to the maintenance of joint integrity in adulthood. Notably, recent genome wide association studies (GWAS) have identified LMX1B as a key genetic risk factor for osteoarthritis, a degenerative joint disease characterised by progressive changes to joint tissues leading to dysfunction.

Despite the clear role that *Lmx1b* plays in limb patterning, the mechanisms through which *Lmx1b* regulates bone and joint formation remain less well understood. Microarray analyses between wildtype (*wt*) and *Lmx1b* knock-out mice have identified several targets involved in skeletal development^4–6^. For example, reduced expression of *Growth differentiation factor 5* (*Gdf5*), a key gene regulator of joint development and a known osteoarthritis risk gene, was observed in *Lmx1b* knockout mice^4^. Similarly, altered expression of proteoglycans (*Keratan, Lumican* and *Decorin*) and extracellular matrix (ECM) genes (*Matrilin-1* and *4)* have been reported^4^. These findings are supported by chromatin immunoprecipitation sequencing (ChIP-seq) against Lmx1b in embryonic mouse limbs, where Lmx1b binding sites in or near potential genes regulated by Lmx1b were identified^7^. Collectively these data suggest a developmental role for Lmx1b in establishing correct skeletal and joint structures.

Importantly, dysregulation of these genes has also been implicated in the pathogenesis of osteoarthritis (OA). OA is characterised by the progressive degeneration of articular cartilage at the joint interface, causing joint inflammation, bone damage and misalignment, pain and reduced quality of life^8^. It is the most common form of chronic joint disease and 14.2% of the global population are estimated to suffer with OA^9^, imposing a significant socioeconomic burden^10^. There are currently no curative treatments available for OA which is in part due its complex pathology and multi-factorial aetiology, involving age, weight, injury and hereditary. Mutations in *Gdf5, Decorin* and *Matrilin-1* and *4* have been associated with OA pathogenesis in human and animal studies^11–14^, while changes in the expression of *Keratan* and *Lumican* can be indicative of OA disease progression^15,16^. Therefore, these data suggest that disruption to pathways regulated by Lmx1b could predispose joints to degeneration in later life.

Recent work has implicated *Lmx1b* in bone homeostasis where *Lmx1b* acts as a negative regulator of the bone morphogenetic protein-2-runt-related transcription factor-2 (Bmp-2-Runx2) signalling cascade; a driver of osteogenesis^17^. Modulation of *Lmx1b* in murine osteoblast precursor cell cultures resulted in dysregulation of key osteogenic markers such as Runx2, alkaline phosphatase (Alpl), bone sialoprotein (Ibsp), and osteocalcin (Bglap), therefore indicating a role for *Lmx1b* in osteoblast differentiation and bone formation^17^. Furthermore, single-cell RNA sequencing from 7-9 week human embryos showed that osteoblast progenitor cells express LMX1B, supporting an evolutionarily conserved role for LMX1B in skeletal regulation^18^.

Beyond early bone skeletal patterning, recent GWAS have linked single-nucleotide polymorphisms (SNPs) in *LMX1B* to hip OA and high bone mass density, highlighting a strong association of LMX1B expression with joint health and maintenance^19–22^. Clinical observations in patients with Nail Patella Syndrome (NPS), caused by mutations in LMX1B, reveal skeletal abnormalities such as scoliosis of the spine, spinal fusions, and premature OA^23^. While it remains unclear whether osteoarthritis in these patients is caused directly by loss of *LMX1B* expression or is secondary to structural changes to the joint, these findings further implicate a role for LMX1B in OA pathogenesis.

Given the role of LMX1B in skeletal patterning and the association of mutations within *LMX1B* with OA^24^, this study looked to explore the effects of loss of *lmx1b* on skeletal maintenance, with a focus on joints, in zebrafish. We have previously shown that of the two *lmx1b* paralogues present in zebrafish (*lmx1ba* and *lmx1bb*), *lmx1ba* is the paralogue primarily associated with the skeletal functions of *lmx1b,* with loss of *lmx1ba* leading to delayed chondrocyte maturation and reduced skeletal growth^25^. Here, we show that skeletal patterning is largely normal, but that *lmx1ba* mutants show progressive skeletal abnormalities from juvenile stages. By 9 months post fertilisation (mpf), adult *lmx1ba* mutants show signs of premature spinal degeneration, intervertebral disc abnormalities and articular cartilage loss in the lower jaw joint; indicative of early-onset OA. We further show changes to cell behaviour that precede pathology. Together these results support a role for LMX1B in skeletal maintenance and demonstrate that disruption of LMX1B function contributes to OA onset and progression.

## RESULTS

### Loss of *lmx1ba* causes premature spinal degeneration

To establish whether *lmx1ba*^−/−^ fish showed signs of premature skeletal degeneration, we assessed micro-computed tomography (µCT) image data from 6-, 9- and 17-month-old wildtype (*wt)* and *lmx1ba* mutants, categorising the location and severity of different spinal abnormalities: intervertebral disc (IVD) calcification, ectopic bone formation and vertebral fusions and misalignments (Figure 1A-D, Supplemental Figure 1A). These spinal abnormalities are associated with IVD degeneration in aged zebrafish^26,27^. *lmx1ba* mutants show high prevalence of abnormalities; with 81.7% of vertebrae displaying at least one abnormality by 17mpf compared to 8% of age matched *wt* vertebrae (Supplemental Figure 1B). At 6mpf, IVD calcification (Figure 1C, *pink arrowhead)* is observed in *lmx1ba* mutants (Figure 1D). By 9mpf, ectopic bone is observed between vertebrae along with vertebral narrowing and misalignments (Figure 1A, *orange* and *green arrowheads;* Figure 1B), and increased IVD calcification. By 17mpf, there is severe disruption to the form and structure of the spine resulting in scoliosis and fusion of vertebrae, along with further ectopic bone deposition and IVD calcification (Figure 1A-C). The severity of spinal degeneration observed in *lmx1ba^−/−^* spines at 17mpf is reminiscent of that observed in much older *wt* fish at 30mpf (Supplemental Figure 2). These data indicate that loss of *lmx1ba* causes a premature and accelerated spinal degeneration phenotype.

**Figure 1.**
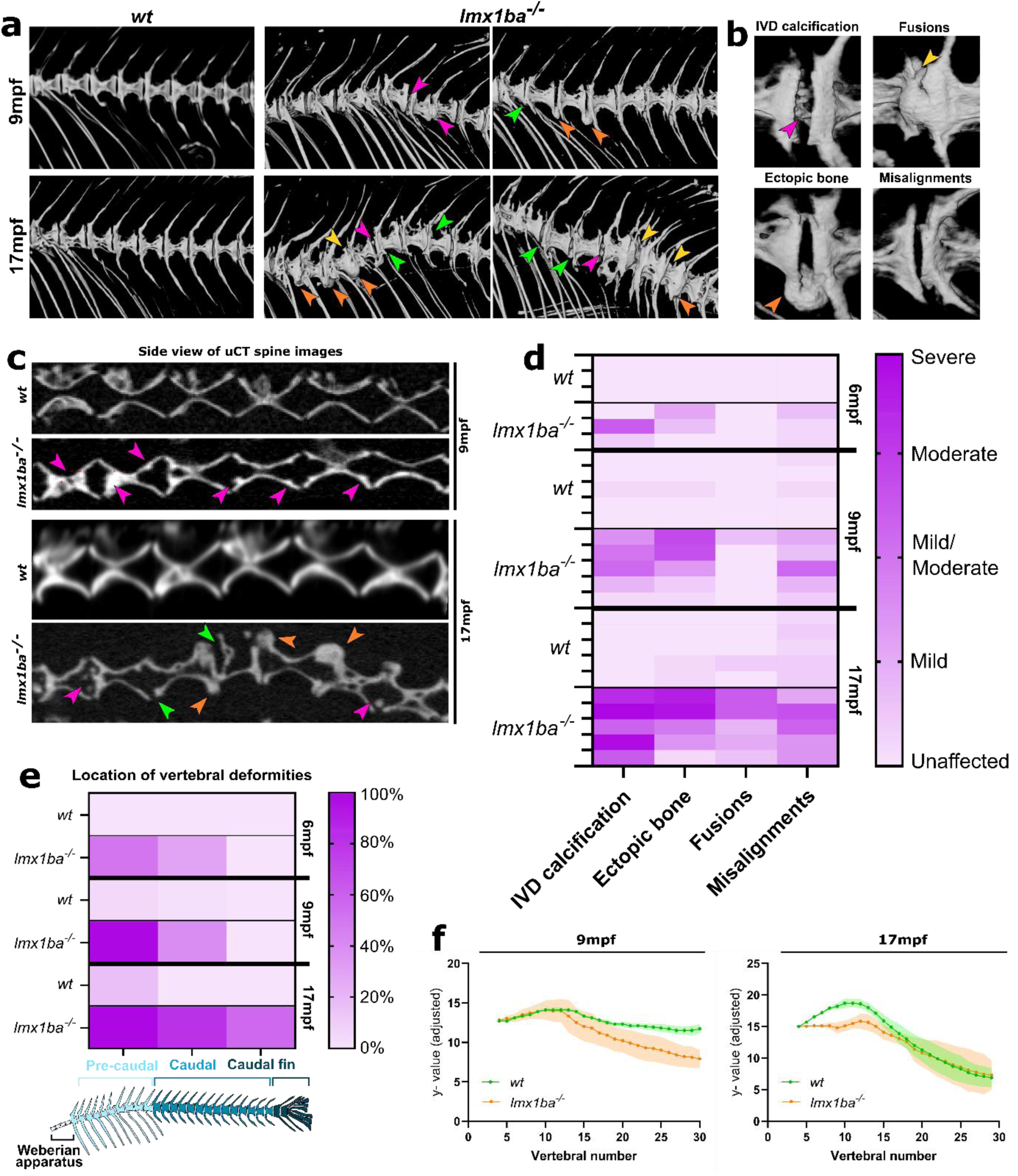
Loss of *lmx1ba* causes progressive spinal abnormalities. **(A)** 3D rendering of µCT images of *wt* and *lmx1ba^−/−^* spines at 9 and 17mpf. Arrowheads show areas of deformation in *lmx1ba^−/−^* spines. **(B)** 3D rendering of µCT images showing spinal ab-normalities common in *lmx1ba^−/−^* fish. **(C)** µCT midline projections of the pre-caudal verte-brae from (A). Arrowheads show regions of IVD calcification (magenta), vertebral misalign-ments (green), bone fusions (yellow) and ectopic bone growth (orange). **(D)** Heat map show-ing spinal morphological changes classified by severity in *wt* and *lmx1ba^−/−^* fish at 6, 9 and 17mpf. N = 3 at 6mpf; N = 5 at 9 and 17mpf. **(E)** Heat map showing the percentage of verte-brae with a deformity in each region of the spine in *wt* and *lmx1ba^−/−^* fish at 6, 9 and 17mpf. N = 3 at 6mpf; N = 5 at 9 and 17mpf. **(F)** Ribbon plot showing overall shape of spine from the pre-caudal to caudal fin vertebrae (from vertebrae 6 to end). N = 3 fish for all ages and geno-types.

To establish whether all regions of the spine are equally affected, we compared pathology in the precaudal, caudal and fin regions of the vertebral volume. At 6mpf, the highest prevalence of degeneration in *lmx1ba^−/−^* fish is in the anterior end of the spine in the pre-caudal vertebrae (from vertebrae 5-14). As the fish age, pathology is increasingly observed in the caudal and caudal fin spine regions, mirroring a similar pattern to *wt* fish, albeit at younger stages (Figure 1D). Given these pathological changes to *lmx1ba^−/−^* spines, we plotted the central position of each vertebral body from the pre-caudal vertebrae to the fin to assess changes to spine curvature (Figure 1E). At 9mpf, *wt* fish show minimal deviation in spine shape between individuals; by contrast, the deviation and displacement of *lmx1ba^−/−^*spines at 9mpf resembles that seen in *wt* fish at 17mpf. By 17mpf, mutants show a curved, scoliotic phenotype (Figure 1F, *orange lines*). Together these data show that spinal phenotypes associated with ageing present much earlier in the *lmx1ba* mutants.

### Osteoblast and osteoclast activity is altered at juvenile stages in *lmx1ba* mutants, indicative of altered remodelling

As spinal fusions and ectopic calcification can be caused by early defects to segmentation in collagen mutants and altered retinoic signalling^28–30^, and as double *lmx1b* knockout zebrafish show defects in notochord inflation^25^, we wanted to establish whether segmentation was normal in the *lmx1ba* mutants. At larval stages, mineralisation of the centrae was slightly delayed at 10 days post fertilisation (dpf) in *lmx1ba^−/−^* fish (average of 6 vertebrae mineralised in *lmx1ba^−/−^* compared to 10 in *wt)* but recovered such that they were indistinguishable from *wt* at 14 and 20dpf. (Supplemental Figure 3A-B). Notochord formation and segmentation also appeared normal (Supplemental Figure 3C-D), indicating that adult changes to spine morphology are not caused by early developmental patterning of the centrae.

At 1mpf, 3D renders of synchrotron radiation-based μCT (SRCT) images of pre-caudal spinal vertebrae showed that the segmentation of *lmx1ba^−/−^* vertebral columns is comparable to *wt* (Figure 2A-B). However, morphological changes in the neural and hemal postzygapophyses were present in *lmx1ba^−/−^* fish where they appear smaller and smoother, suggestive of increased remodelling compared to *wt* (Figure 2B, *arrowheads*). Neural and hemal postzygapophyses form at the posterior ends of centra at the dorsal and ventral sides respectively, where ligaments associated with spine stability attach the bone^31^.

**Figure 2.**
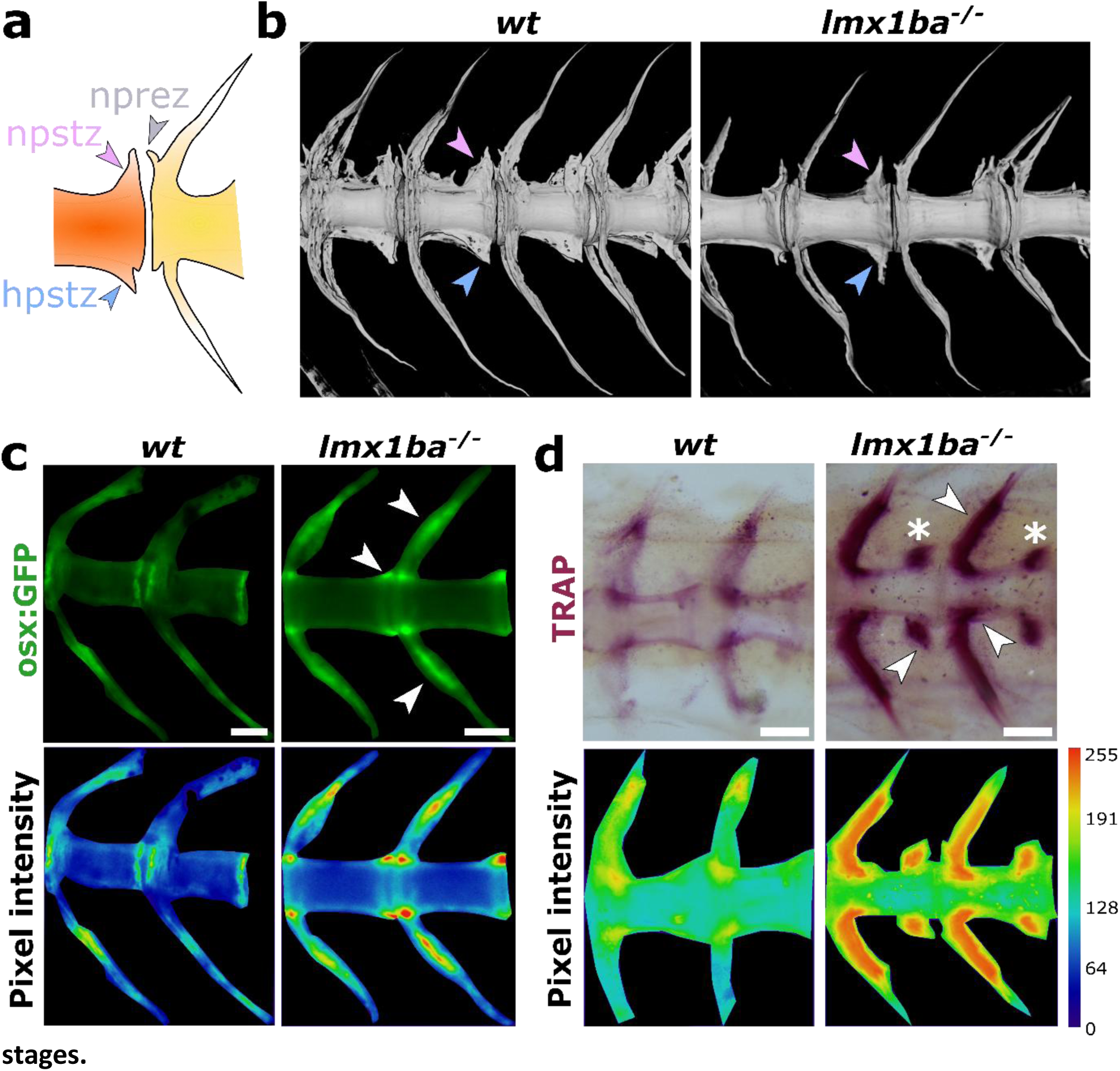
Increased osteoblast and osteoclast activity in *lmx1ba* mutants at juvenile stages. **(A)** Schematic showing location of neural postzygapophyses (npstz), hemal postzygapophyses (hpstz) and neural prezygapophyses (nprez) in a 1mpf spine. Ligaments from npstz attach to nprez for spinal stability. **(B)** 3D volumetric rendering of synchrotron radiation-based μCT images of *wt* and *lmx1ba^−/−^*spines at 1mpf. Arrowheads show morphological differences in npstz (pink) and hpstz (blue) formation between *wt* and *lmx1ba^−/−^* spines. N = 3 per group. Stereomicroscope images of *wt* and *lmx1ba^−/−^* **(C)** *osx-nls:eGFP* spines and **(D)** TRAP-stained spines at 25dpf. Pictures were processed to show pixel intensity (blue=low intensity, red=high intensity), to visualise differences in osteoblast or osteoclast activity, respectively. White arrowheads show regions of increased osteoblast or osteoclast expression, respectively, asterisks show npstz. Scale bars = 100µm. N = 8 per group for all.

As we observed changes to these connective tissue-associated structures, and as *in vitro* studies in murine osteoblast/clast precursor cells have indicated a role for *Lmx1b* as a negative regulator of both osteoblast and osteoclast differentiation^17,32^, we wanted to explore the effect of loss of *lmx1ba* on osteoblasts and osteoclasts. *lmx1ba* mutants showed increased *osterix* expression in ends of the vertebra and in the neural arches compared to *wt* fish at 25dpf (Figure 2C, *white arrowheads*). Tartrate-Resistant Acid Phosphatase (TRAP) staining, used to identify osteoclast activity, was also more intense in the arches and neural postzygopophyses in *lmx1ba*^−/−^ spines (Figure 2D, *white arrowheads and asterisk,* respectively), suggestive of higher remodelling activity. These minor morphological alterations to the zygopophyses are maintained at 3mpf, whilst histological analysis of spinal sections showed early vertebral bone misalignments in the *lmx1ba* mutants (Supplemental Figure 4). Together these data suggest that loss of *lmx1ba* does not alter developmental spinal segmentation but does lead to altered remodelling that could underpin the progressive pathology observed.

### Loss of *lmx1ba^−/−^* results in IVD degeneration involving increased fibrosis, collagen disruption and disc herniation

As *lmx1ba* mutants showed changes close to the tissue interface between the bone and the IVD, we wanted to next establish if there were changes to the IVD in the mutants. IVDs connect adjacent vertebrae forming a fibrocartilaginous joint that allows for flexible spine movement as well as shock absorption. In mammals, IVDs consist of an elastic and hydrated gel-like centre termed the nucleus pulposus (NP), which is surrounded by an outer fibrous ring of connective tissue termed the annulus fibrosus (AF). Recently, several papers have identified NP, AF and endplate structures in zebrafish with similar cellular properties and organisation to those of mammals^26,27,33^. During ageing or in disease states, zebrafish models show changes to NP and AF architecture with strong resemblance to IVD degeneration in mammals, including humans^26^.

Histological analysis of adult *wt* and *lmx1ba^−/−^*zebrafish spines at 9mpf and 17mpf was performed, using toluidine blue and keratin staining on sequential sagittal sections. At 9mpf, accumulation of fibrotic tissue and vertebral misalignments were evident in some *lmx1ba^−/−^*IVDs (Figure 3A, *yellow and green arrowheads*), whilst others had a milder phenotype more comparable to *wt* at 17mpf, showing only initial disorganisation of vacuolated cells at the NP ends (Figure 3A, *red arrowhead*). By 17mpf however, *lmx1ba^−/−^* fish showed complete disorganisation of vacuolated cells and fibrosis of the NP, resulting in herniation of the disc in some regions (Figure 3A, *yellow arrowheads* and *white asterisks*). In adult zebrafish, the central septum of the NP is formed mainly of keratin and collagen type I^26^. These components can be visualised using pan-keratin staining where keratin is labelled in red/orange, collagens (including collagen type I) in yellow, and glycosaminoglycans in blue (Figure 3C). At 9mpf, *lmx1ba^−/−^* fish show a similar phenotype to older 17mpf *wt* fish, with a slight increase in keratin staining of the NP septum (Figure 3B). At 17mpf, there is an accumulation of keratin in the *lmx1ba^−/−^* NP indicating fibrotic scarring of the IVD (Figure 3B, *yellow arrowhead),* as typically seen in much older fish^26^.

**Figure 3.**
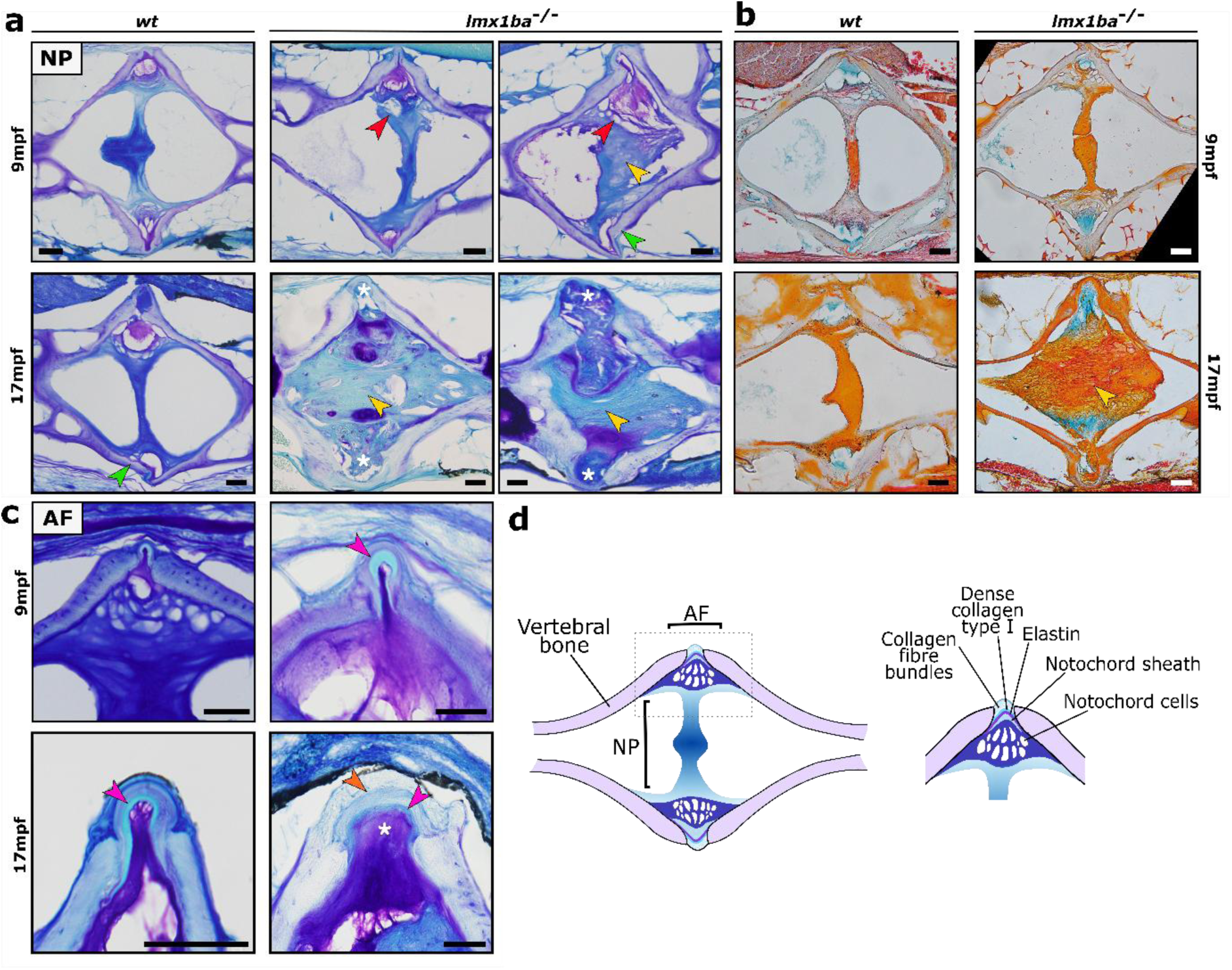
IVDs show premature degeneration and scarring in *lmx1ba* mutants by histolog-ical analysis. Histological sections of adult *wt* and *lmx1ba^−/−^* IVDs at 9mpf and 17mpf stained with **(A)** Toluidine blue or **(B)** keratin for the nucleus pulposus (NP), or **(C)** Toluidine blue for the annulus fibrosus (AF). Arrowheads show forms of degeneration; disorganisation of vacuolated cells (red), fibrosis (yellow), vertebral misalignment (green), thickened elastin layer in AF (magenta), abnormal collagen layer in AF (orange) or herniation of IVD (white asterisks). Scale bars = 50µm. **(D)** Schematic showing structure and layout of NP and AF in zebrafish. N = 3 for *wt* 9mpf and *lmx1ba^−/−^* 17mpf, N = 6 for *lmx1ba^−/−^* 9mpf and N = 5 for *wt* 17mpf.

Analysis of the AF region showed evidence of a thickened collagen type I layer in 9mpf *lmx1ba^−/−^* fish, resembling the phenotype of older *wt* AF (Figure 3C, *magenta arrowheads)*. At 17mpf, the collagen fibre layer appears thicker and more disorganised, indicative of fibrosis and NP degeneration^27,34^, and herniation of the disc is apparent (Figure 3C, *orange arrowhead* and *white asterisk)*. In conclusion, loss of *lmx1ba* leads to premature degeneration of the IVD involving accumulation of fibrotic tissue in the NP, thickening of collagen layers in the AF and ultimately disc herniation.

### Proteoglycan and glycosaminoglycan profiles are altered in the spines of *lmx1ba* mutants

Proteoglycans (PGs), such as aggrecan, versican and lubricin are found in most matrix rich tissues and are core components in cartilage. Lubricin is a mucinous glycoprotein found in synovial fluid and on the surface of cartilage^35^ and is essential for joint homeostasis, with a reduction in lubricin expression leading to rapid cartilage damage^36^. PGs are comprised of glycosaminoglycan (GAGs) chain(s) covalently bound to a protein core. GAGs are long negatively charged polysaccharides which provide essential hydration, structural integrity and shock absorption for cartilage and are also involved in cell signaling^37,38^.

GAGs are classified into four groups based on their core disaccharide structures and include: heparan sulphate (HS), chondroitin sulphate, (CS), keratan sulphate (KS) and hyaluronic acid (HA). Chondroitin sulphate is the most common GAG in cartilage composing roughly 80% of total GAGs^39^ and is the major type in mineralized bone matrix where it supports bone stiffness and dissipation of energy^40,41^. All GAGs, apart from HA, may be sulphated, altering their composition and function. Changes in sulphation influence macrostructure and binding and hence, changes in sulphation can be inferred as changes in function^42^. For CS, there are 4 main categories of CS based on the sulphation position(s) on their glucuronic acid (GlcA) and N-acetylgalactosamine (GalNAc) residues. These are known as CSA and CSC (which are mono-sulphated) and CSB, CSD and CSE (which are di-sulphated), with CSA and CSC being the two main forms found in cartilage and bone tissue^43,44^. Alterations to CS levels and sulphation are common hallmarks in OA^45,46^; polymorphisms in genes regulating chondroitin sulphation and synthesis such as CHST11 have been linked to increased OA risk and CHSY1 to disc degeneration^47,48^.

To analyse the GAG composition of *wt* and *lmx1ba^−/−^*spines, trapped ion mobility spectrometry mass spectrometry imaging (TIMS-MSI) was used as previously described^49^. Chondroitin sulphate (CS) analysis revealed marked changes in CS composition and localisation (Figure 4A-D). In *wt* fish, CSA was seen in the vertebral bone and IVD, however, CSC was localised to the AF of the IVD. In *lmx1ba^−/−^* fish, CSC was localised instead to the vertebral bone and not the IVD or AF (Figure 4A-B).

**Figure 4.**
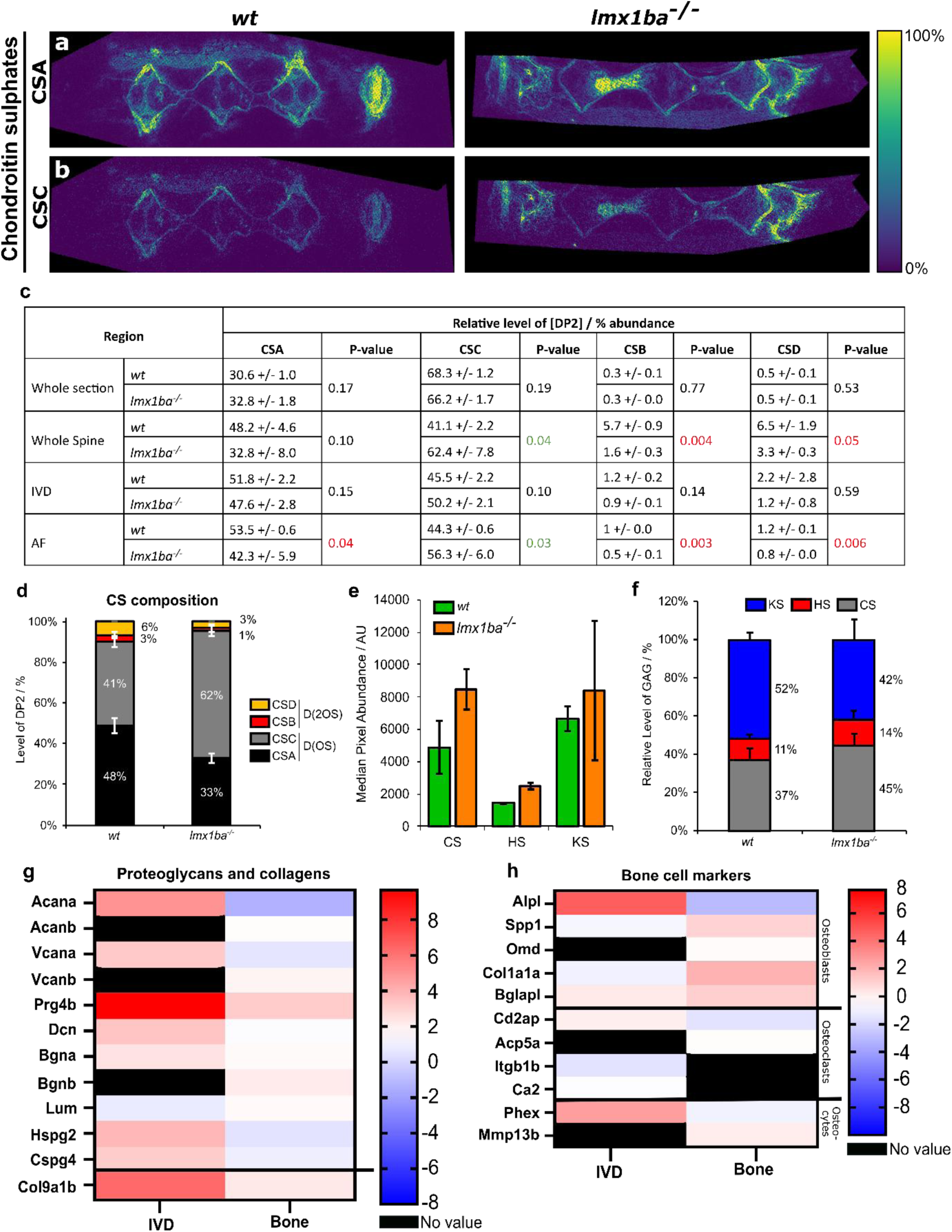
GAG and proteoglycan composition is altered in IVDs of *lmx1ba* mutants. Trapped ion mobility spectrometry mass spectrometry imaging (TIMS-MSI) of chondroitin sulphates (CS) DP2s comparing **(A)** CSA vs **(B)** CSC levels between for *wt* and *lmx1ba^−/−^* fish at 9mpf. **(C)** Relative abundance of detected CS disaccharides. Students T-test performed be-tween *wt* and *lmx1ba^−/−^*fish of the same region. Green text = significant increase in CS in *lmx1ba^−/−^*compared to *wt*; red text = significant decrease in CS in *lmx1ba^−/−^*compared to *wt.* **(D)** Compositional analysis of CS levels by DP2. Compositional analysis of GAGs level by **(E)** median abundance and **(F)** relative level. CS = chondroitin sulphate; HS = heparan sulphate; KS = keratan sulphate. N = 2 per group. Plot of log2 of fold change in abundance of **(G)** proteoglycans and collagen proteins and **(H)** bone cell markers relative to *wt* for IVD and bone tissue, respectively from mass spectrometry. N = 1 per group. Black cells indicate where one group did not have a value for that protein and they were excluded.

To understand further the change in CS sulphate patterns, we performed spatially resolved disaccharide analysis. TIMS profiles were extracted and analyzed as described in^50^ to yield pseudo-disaccharide profiles (Figure 4C-D). Analysis of the whole spines showed significant changes in CSC (P=0.04), CSB (P=0.004) and CSD (P=0.05) levels in *lmx1ba^−/−^*, indicating a localised change to CS levels. In particular, dysregulation to CS profiles was observed in the AF region (Figure 4C) where levels of CSA (P=0.04), CSC (P=0.03), CSB (P=0.003) and CSD (P=0.006) were all dramatically altered in *lmx1ba* mutants.

Finally, we examined overall GAG composition and abundance in the tissue. Although increases in heparan sulphate (HS) and keratan sulphate (KS) levels were seen in *lmx1ba^−/−^*spines, quantitative analysis of HS disaccharides and KS monosaccharides revealed no significant changes (Supplemental Figure 5). CS levels, however, did show increased intensity (Figure 4E) as confirmed by the comparison of relative GAG levels in the spine (Figure 4F). Interestingly, increases to CS along with altered location have previously been shown in aged zebrafish spines^51^. Together, these data indicate that a change in CS, both by composition and by abundance, is present in *lmx1ba^−/−^* fish, especially in the AF region of the IVD.

To identify protein changes between *lmx1ba^−/−^* and *wt* spines we used laser capture microdissection (LCM) to isolate both IVD and vertebral bone regions for MS analysis (Supplemental Figure 6). Overall, 1138 proteins for vertebral bone and 1715 proteins for IVD were quantified between *wt* and *lmx1ba^−/−^* (Supplemental Dataset 1) and the abundances of proteins across each tissue were normalised to *wt*. Of the 11 PGs identified, all but one (lumican) showed increased abundance in *lmx1ba^−/−^* IVD but were decreased in mutant bone compared to *wt* (Figure 4G). This aligns with the GAG analysis where CS levels (a core component of PGs, especially aggrecan, versican, decorin and biglycan) are increased in *lmx1ba^−/−^*spines, further indicating that loss of *lmx1ba* alters PG and GAG composition within the spine. Increases in osteoblast cell markers were observed in *lmx1ba^−/−^* vertebral bone, in line with the elevated *osx:GFP* expression at 1mpf, although osteoclast markers showed no significant changes (Figure 4H). Interestingly, in *lmx1ba^−/−^* IVDs, proteins involved in bone mineralisation such as *Alpl*, a key bone mineralisation enzyme, and *Phex,* an osteocyte marker, showed a 5- and 3- fold increase in abundance, respectively, compared to *wt* (Figure 4H). This is likely driving the increased IVD calcification seen in *lmx1ba^−/−^*at this age, demonstrating that *lmx1ba* is important for maintaining bone cell homeostasis.

### Loss of *lmx1ba* leads to heterogeneity in bone mineral density and increased bone fragility

Given these molecular changes to bone and ECM composition and the changes to osteoblast and osteoclast behaviour, we wanted to test the functional relevance of these changes at a tissue level. Whole spines were segmented from µCT images and bone mineral density (BMD) calculated for *wt* and *lmx1ba^−/−^* at 9mpf (Figure 5A-B). BMD was elevated in *lmx1ba^−/−^*spines (Figure 5B). The standard deviation in BMD measurements was also significantly increased, suggesting a high mineral heterogeneity across *lmx1ba^−/−^* spines (Figure 5C); an indicator of reduced bone quality and fracture risk often associated with aged spines^26,52^.

**Figure 5.**
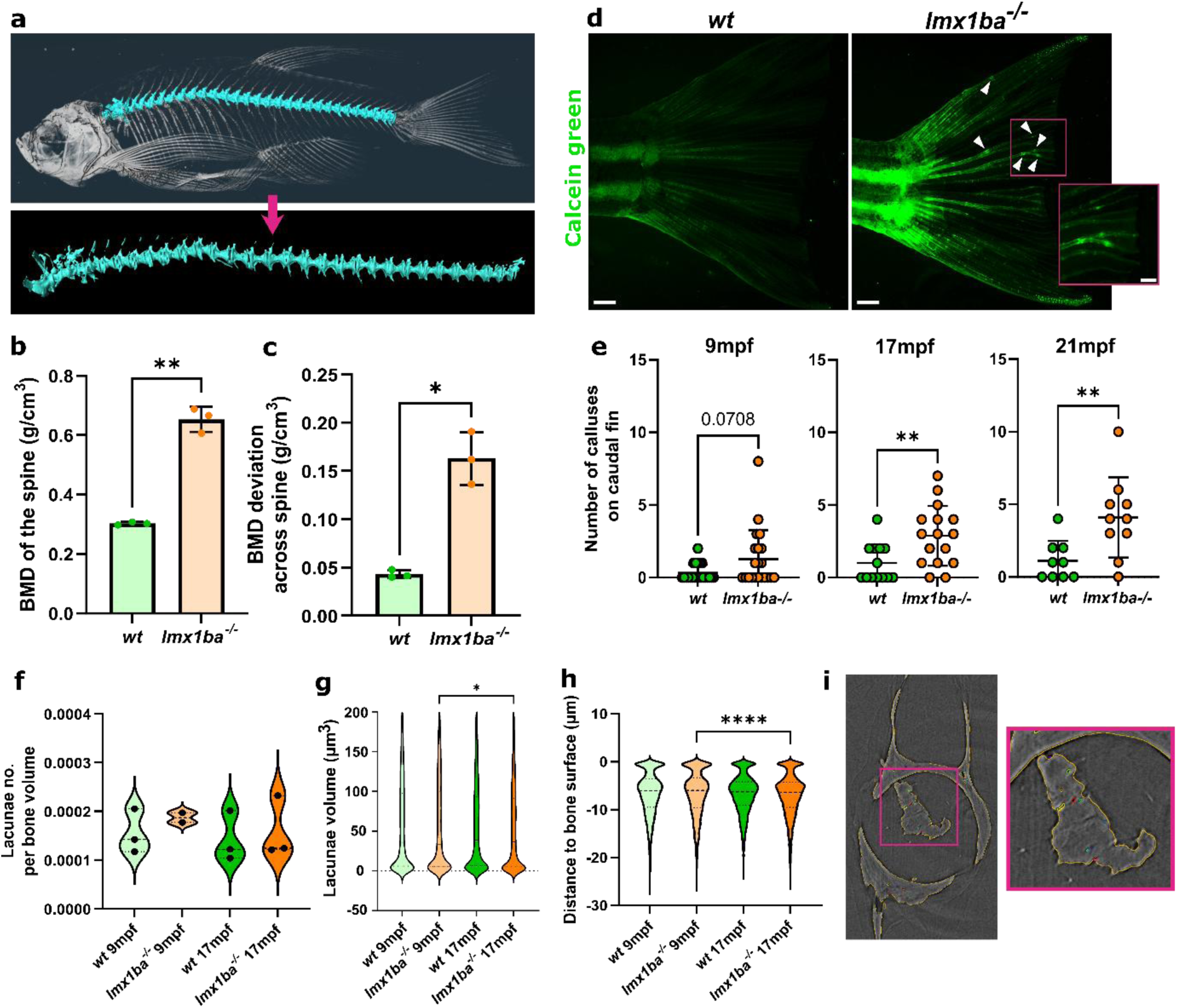
Bone density and fragility is altered in *lmx1ba* mutants but this is not due to changes to osteocyte patterning. **(A)** 3D volume rendering of µCT image showing segmentation of whole spine from an adult *wt* fish for bone mineral density (BMD) analysis. Graphs showing **(B)** volumetric BMD and **(C)** BMD deviation across the whole spine from *wt* and *lmx1ba^−/−^* fish at 9mpf. N = 3 per group. Welch’s T-test performed for both where \*\**P=*.0043 and \**P=*.0153. **(D)** Stereomicroscope images of adult caudal fins live stained with calcein green to visualise bone calluses as shown by white arrowheads. Scale bars = 500 µm and 200 µm for inset. **(E)** Quantification of number of endogenous (i.e. not experimentally caused) calluses in caudal fin rays at 9mpf (N = 26 for *wt* and 25 for *lmx1ba^−/−^),* 17mpf (N = 13 for *wt* and 16 for *lmx1ba^−/−^*) and 21mpf (N = 9 for *wt* and 10 for *lmx1ba^−/−^*). Mann-Whitney T-test performed at 9 and 17mpf where *\*\*P=*.0083, and Welch’s T-test performed at 21mpf where *\*\*P=*.0094. **(F)** Violin plot of num-ber of lacunae per bone volume. **(G)** Violin plot of lacunae volume. One-way ANOVA per-formed where *P=*.0198. **(H)** Violin plot of lacunae distances from the bone surface. One-way ANOVA performed where *P= <*.0001. N = 3 for all apart from *lmx1ba ^−/−^* 9mpf where N = 2. **(I)** Z-slice from SRCT image of *lmx1ba*^−/−^ IVD at 17mpf following lacunae analysis workflow. Pink inset shows zoom of ectopic bone in IVD; lacunae highlighted by coloured outlines, bone highlighted by yellow outline.

Bone fragility was assessed by counting the number of calluses found in caudal fin ray bones as an indicator of spontaneous fracture occurrence^53^ (Figure 5D). From 9mpf, *lmx1ba* mutants showed an increase in average callus number which was significant by 17mpf (Figure 5E), suggesting that loss of *lmx1ba* leads to increased bone fragility.

The coordination of bone remodelling is largely regulated by osteocytes: the most abundant bone cells, located in cavities within the bone matrix known as lacunae and interconnected via cell extensions known as canniculae. Through this network, osteocytes can sense and respond to changes in mechanical loading or bone damage and signal to osteoblasts and osteoclasts. Variation in osteocyte morphology and location have been observed in ageing or disease states in humans and zebrafish and can indicate impaired mechanosensory ability and perturbed bone homeostasis^26,54–56^.

Using an automated workflow, lacunae were detected and analysed from spinal regions of *wt* and *lmx1ba^−/−^* fish from SRCT images at 9- and 17mpf. No significant changes to lacunae number, volume or depth were seen between *wt* and *lmx1ba^−/−^* spines (Figure 5F-H), suggesting that osteocyte formation and patterning in the cortical bone is unaffected in *lmx1ba* mutants, consistent with our observations that early spinal patterning is largely normal. Interestingly, lacunae were also detected within regions of IVD calcification found in *lmx1ba^−/−^*spines (Figure 5I), indicating that these deposits are ectopic, ossified bone. Together with the disparity in BMD across the spine, osteocytes in these ectopic bone regions may be influencing local remodelling activity, contributing to the progressive phenotype observed in *lmx1ba* mutants.

Alterations to BMD and bone fragility can impact functional performance of the skeletal system. To assess the functional impact of these changes in the *lmx1ba* mutants at an organismal level, spine curvature during swimming was measured. Observing the three greatest right and left movements and the average right and left curvature for each fish at 9mpf (Figure 6), *lmx1ba* mutants showed increased curvature compared to *wt* suggesting greater spinal flexion and reduced spinal stability. This demonstrates that loss of *lmx1ba^−/−^*affects the structural integrity of the spine leading to mechanical and functional changes to body movement.

**Figure 6.**
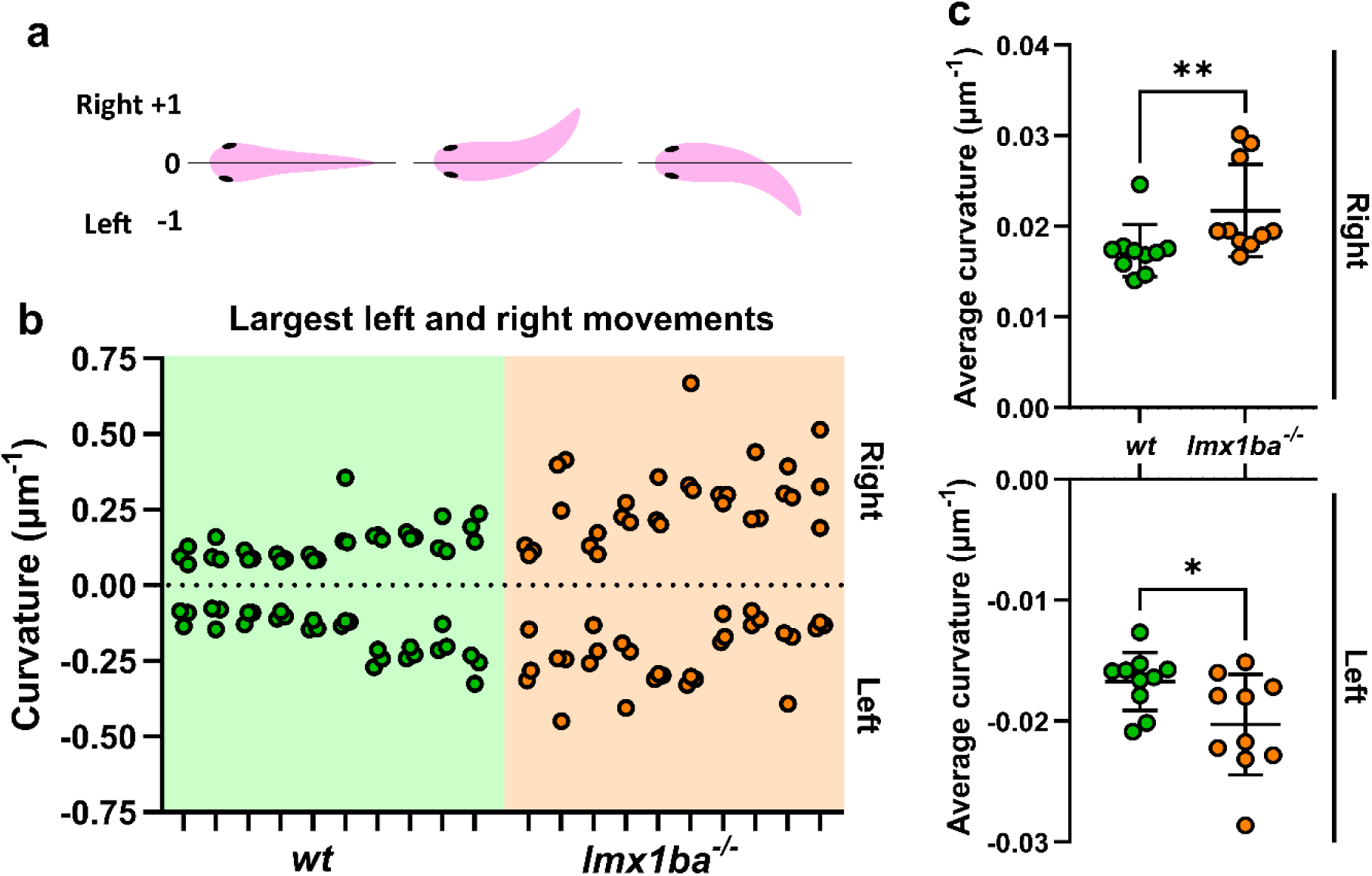
Bone mechanics is altered in *lmx1ba* mutants. **(A)** Schematic showing body curvature measurement by the modular image analysis (MIA) program. When fish is straight = 0; when fish bends right = up to +1; when fish bends left = up to −1. Measurements taken per frame of video. Graphs showing **(B)** largest 3 left and right movements per *wt* or *lmx1ba^−/−^* fish at 9mpf. Each dash on x-axis indicates measurements from 1 fish. And **(C)** average spine curvature to right or left per fish. Mann-Whitney and Welch’s T-test performed for right and left measurements, respectively where \*\**P=*.0039 and *\*P=*.034. N = 10 for both groups.

### Skeletal abnormalities caused by loss of *lmx1ba* are observed in other mobile joints, while immobile joints are preserved

To assess whether the bone and joint changes observed in the spine are reflected across other skeletal elements, we wanted to establish whether progressive degeneration was observed in other joints. We therefore compared the jaw joint, a synovial joint that forms early in zebrafish development and cranial sutures which are immobile fibrous joints. At 9mpf, histological analysis of *lmx1ba^−/−^* jaw joints showed increased joint widening and premature articular cartilage degeneration at the joint site compared to *wt* (Figure 7A). This is confirmed by 3D rendering of SRCT images of the lower jaw joint where misalignment of the joint is also observed in the *lmx1ba* mutants (Figure 7B, *yellow arrowhead*). However, analysis of cranial sutures, the fibrous joints between skull bones considered static by adulthood, showed no abnormalities (Figure 7C-E). This indicates that the skeletal changes triggered by loss of *lmx1ba* are generalisable to other mobile joints, but not to static joints, highlighting a selective requirement for *lmx1ba* in mechanically active joints. Together our findings demonstrate a role for *lmx1ba* in the maintenance of skeletal tissue integrity across the whole joint unit, loss of which leads to progressive development of osteoarthritic pathology.

**Figure 7.**
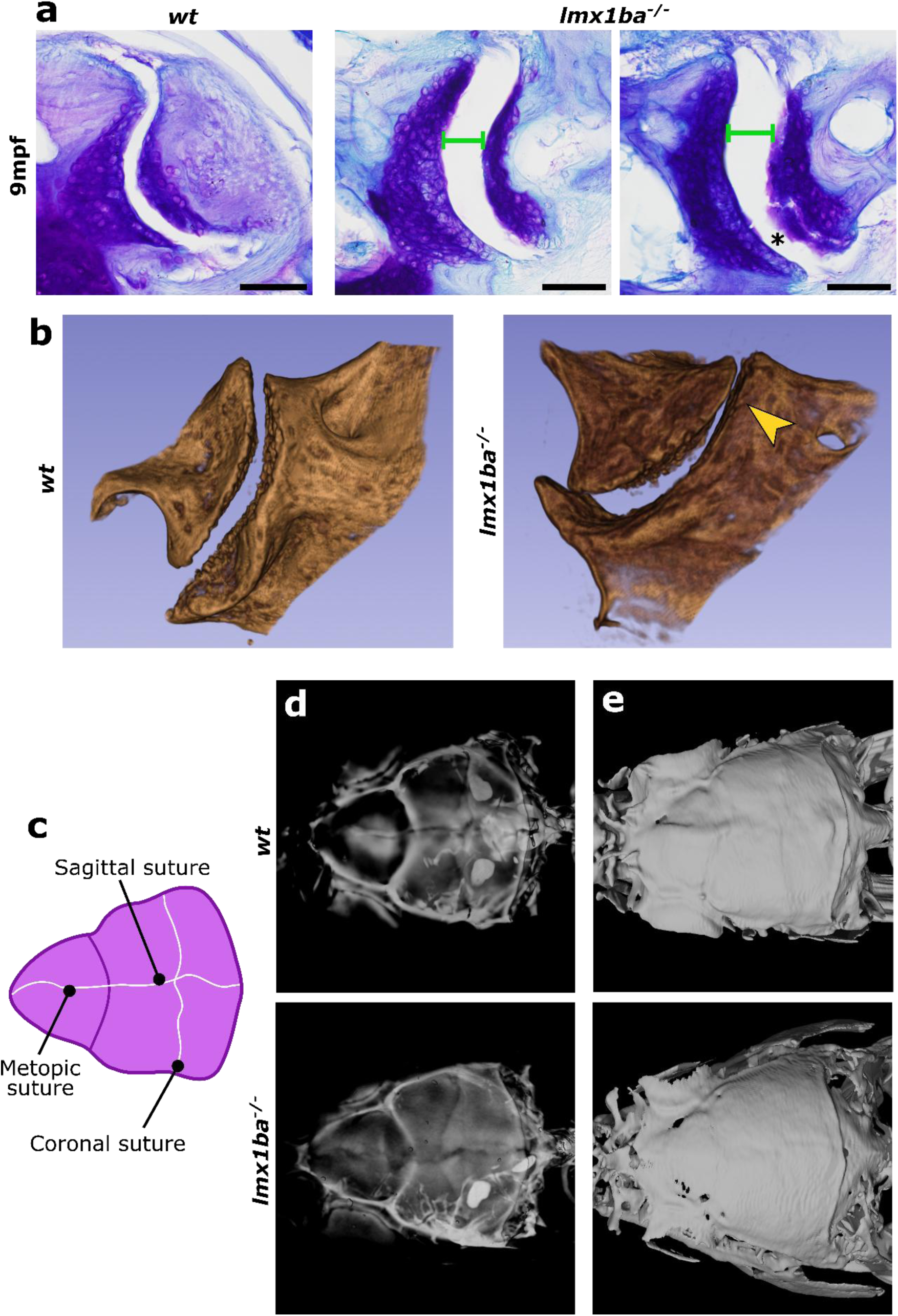
Effect of *lmx1ba* mutation is restricted to mobile joint elements. **(A)** Histological sections of adult zebrafish lower jaw joints at 9mpf stained with Toluidine blue. Green line indicates wider gap in jaw joint; asterisk shows articular cartilage degenera-tion. Scale bars = 500 µm. N = 3 per group. **(B)** 3D volumetric rendering of synchrotron radia-tion-based μCT images of *wt* and *lmx1ba^−/−^* lower jaw joints at 9mpf. Yellow arrowhead shows misalignment of jaw joint in *lmx1ba* mutants. N = 3 per group. **(C)** Schematic showing position of cranial sutures in zebrafish skulls. **(D)** Volume rendering and **(E)** segmentation of 9mpf *wt* and *lmx1ba^−/−^* skulls from µCT images. N = 5 per group.

## DISCUSSION

Our findings identify LMX1B as a critical regulator of skeletal homeostasis whose dysregulation drives progressive osteoarthritis through disruption of bone–cartilage–disc integration. Notably, skeletal patterning in *lmx1ba* mutants is largely preserved during early development, with craniofacial and axial structures forming without overt morphological defects. Consistent with this, osteocyte lacunar organisation is unchanged, indicating that osteocyte embedding is established appropriately during the development of the vertebral centrae, with pathological changes to both bone and disc occurring later. This temporal separation between normal developmental patterning and later-onset degeneration reveals a key role for Lmx1ba in maintenance, rather than establishment, of skeletal integrity. Thus, this supports the concept that osteoarthritis can arise from failure of long-term tissue homeostasis, where initially subtle changes to the joint become amplified over time through dysregulation of the whole joint unit leading to altered cell behaviour.

We observe that OA-like pathology upon loss of *lmx1ba* emerges by early disruption of bone remodelling dynamics. Localised changes in osteoblast and osteoclast activity are detectable from 25dpf, focused in regions where tissues integrate, preceding overt structural pathology by several months. This suggests impaired coupling between bone formation and resorption, a process essential for maintaining skeletal quality and joint stability. Increased bone mineral density (BMD), together with pronounced heterogeneity of BMD, indicates altered mineral distribution and reduced bone quality, features associated with osteoarthritis and subchondral sclerosis^57^.

Importantly, we also observe an increased incidence of spontaneous fractures in the fin skeleton, supporting the idea that heterogeneity in mineralisation creates localised foci of mechanical weakness. We speculate that the juxtaposition of regions of high and low BMD generates stress concentrations that predispose to microdamage and fracture. In this context, microinjury, or localised points of aberrant strain, could represent an early and underappreciated driver of disease progression, whereby repeated cycles of damage and repair contribute to aberrant remodelling and tissue degeneration. Indeed, alterations to local density properties have been linked to fracture risk in both zebrafish and humans^53^.

Concomitant with these changes in bone, we observe premature degeneration of the intervertebral disc (IVD) characterised by disruption of glycosaminoglycan and proteoglycan content. Rather than indicating a uniform shift in matrix composition, our data support a model of disrupted and spatially heterogeneous ECM, in which local changes to GAG and proteoglycan abundance lead to altered mechanical behaviour. Under conditions of cyclic loading this heterogeneity in the IVD and AF is likely to amplify stresses over time loading to progressive structural failure. The parallel disruption of bone remodelling and disc composition points to a breakdown in coordinated regulation across the bone–cartilage–disc axis, supporting the concept of osteoarthritis as a whole-joint disease^58^.

Our data further suggest that biomechanical dysfunction is both a consequence and a driver of disease progression. Early alterations in the vertebral arches and zygapophyses are likely to influence joint mobility, and indeed we observe increased spinal flexion during swimming, indicative of altered mechanical performance at the organismal level. We propose that these initial changes in joint mechanics, compounded by localised bone weakness and susceptibility to microinjury, initiate a feed-forward cycle in which altered load distribution exacerbates remodelling defects and accelerates disc degeneration. The later emergence of ectopic ossification, vertebral fusion, and misalignment is consistent with progressive mechanical instability and compensatory stiffening. Interestingly, changes to collagen 11 and to disc integrity also led to altered swim behaviour and to spinal pathology^59^. Similar relationships between altered biomechanics, bone adaptation, and cartilage degeneration are well established in osteoarthritis, with subtle changes to joint morphology predicting later disease^60,61^, supporting the relevance of this model.

Jaw joint pathology in the mutants suggests that this mechanism is generalisable across mobile joints. Increased joint space and premature cartilage degeneration in the jaw indicate that Lmx1ba-dependent regulation of joint integrity is not restricted to the axial skeleton but operates across multiple joint types. This multi-site involvement is consistent with systemic disruption of joint homeostasis and suggests that Lmx1ba functions as a global regulator of skeletal tissue balance. Interestingly we do not observe obvious morphological changes and ectopic bone in the cranial sutures, further suggesting that progression to pathology is exacerbated by mechanical disruption in joints. At a functional level, the preservation of early skeletal architecture alongside defects in remodelling dynamics supports a model in which *lmx1ba* is required to maintain coordinated activity between skeletal cell populations and tissues over time. Loss of this coordination results in progressive uncoupling of bone with surrounding softer connective tissues of the articular cartilage in the jaw and the intervertebral disc, ultimately leading to structural failure.

Together, our study demonstrates that disruption of *lmx1ba* does not impair initial skeletal formation but rather compromises long-term maintenance, leading to progressive osteoarthritis through coordinated failure of homeostasis in the boundaries where bone and soft tissues integrate. By linking early remodelling defects to biomechanical dysfunction and structural degeneration, we define a temporal framework for disease progression. These findings support a unifying principle in which sustained activity of developmental regulators is required to preserve adult joint function, and that their dysregulation represents an initiating event in osteoarthritis. More broadly, this work highlights the interplay between mechanical and biological processes as a central axis of disease and identifies *lmx1ba* as a key regulator of skeletal tissue balance.

## MATERIALS AND METHODS

### Zebrafish husbandry and lines used

Zebrafish (*Danio rerio*) were raised and maintained under standard conditions^62^. Experiments were approved by the local ethics committee (Animal Welfare and Ethical Review Body of the University of Bristol) and performed under UK Home Office project licence (PP4700996). *lmx1ba ^bsl29620^* mutants (previously generated and described by us) have a 7 bp deletion in the conserved homeobox domain causing a premature stop codon at amino acid 219^25^. Transgenic lines used: *Tg(Ola. Sp7:nlsGFP)^zf132^* for osteoblasts^63^, Tg(*kita:Gal4;UAS:mCherry*) for vacuolated notochord cells^64^ and *Tg(col9a2:GFPCaaX)* for the alpha-2 subunit of collagen type IX^65^.

### Micro-Computed tomography (µCT)

Adult *wt* and *lmx1ba^−/−^* fish were culled and fixed in 4% PFA for 7 days and sequentially dehydrated to 70% ethanol. Fish were scanned using a Nikon XT H225ST computed tomography (CT) scanner (Nikon, Japan) with a voxel size of 21 μm (scan settings 130 kV, 150 µA, 0.5-s exposure, 3141 projections). Images reconstructed using CT Pro 3D software (Nikon). “Phantom” samples of known hydroxyapatite concentrations (0.25 and 0.75 g/cm^−3^ calcium hydroxyapatite) were scanned using identical settings to allow conversion of greyscale attenuation values into physical density units. Avizo (v2025.1, Thermo Fisher Scientific) used for 3D volume rendering, segmentations and image acquisition.

### µCT segmentation and bone mineral density (BMD) quantification

CTPro3D was used to calibrate µCT scans by sampling the phantom across multiple slices to derive a mean scaling factor which was applied to all scans to ensure consistent density normalisation. Using Avizo, a consistent threshold range was applied across all specimens to isolate mineralised tissue. Threshold-based segmentation identified bone voxels and anatomical boundaries were refined manually. For each specimen, the vertebral spines were manually segmented and ribs and associated processes removed. Surfaces were generated from label fields and material statistics were computed for each labelled region, yielding voxel-based summary metrics including mean greyscale intensity. BMD was calculated by multiplying the mean voxel intensity values by the calibrated density-per-pixel scaling coefficient derived during phantom calibration to convert these values to density (g/cm⁻³).

### Synchrotron radiation-based μCT (SRCT)

*wt* and *lmx1ba^−/−^* fish at 1, 2, 3, 9 and 17 mpf (*n* = 3 for each group) were scanned at the I13-1 beamline of the Diamond Light Source (DLS) using a pink beam of approximately 27 KeV mean X-ray energy, an isotropic voxel size of 0.33–0.68 μm, 1501 projections, and 50 ms exposure time. Reconstructions were processed using single propagation distance phase retrieval with a *β/δ* ratio of 2.0. Reconstructions were analysed using Avizo.

### Stereomicroscope imaging of zebrafish

Live zebrafish at larval and adult stages were imaged using a DFC700T camera mounted to a Leica MZ10F modular stereomicroscope system (Leica Microsystems) at 1-12x magnification. For live imaging, fish were anaesthetised using 0.1 mg/ml MS222 (Tricaine methane sulfonate) diluted in Danieaus and imaged laterally.

### Live bone staining

Live larval and adult fish were incubated for 1 hour in either 0.1 % Alizarin Red S (Sigma, A5533) or 40μM calcein powder (pH 8.0) (Sigma-Aldrich, C0875) in Danieau solution, rinsed twice and imaged as above.

### Tartrate-resistant acid phosphatase staining

Juvenile fish at 25dpf were culled and fixed in 4% PFA overnight before staining in 2ml TRAP solution (0.2693 mM Naphthol AS-MX phosphate in formamide, 1.6 mM Fast Red Violet LB salt in 50 mM sodium L-tartrate in 0.1M sodium acetate buffer (pH 5)) for 6 hours in the dark at room temperature on a rocker. Samples were washed in PBS-Tween (0.1%) three times and re-fixed with 4% PFA for 30 minutes. Samples were bleached in 0.5% KOH and 3% H2O2 for 25 minutes, washed and the spines dissected out. Spines were cleared in 10% glycerol and 0.5% KOH for 24 hours before stereomicroscope imaging.

### Histology

*wt* and *lmx1ba^−/−^* fish at 1-17mpf were fixed in 4% PFA, decalcified in 50 mM EDTA and dehydrated before embedding in Technovit. Lateral serial sections (2–4 µm) were prepared using a Prosan HM360 microtome. Slides were dewaxed and stained in either 0.04% Toluidine Blue in 0.1M sodium acetate for 15 minutes or a pan-keratin stain as described previously^66^, then washed in increasing serial ethanol washes, dried and mounted using DPX. Images acquired on an Olympus BX53 fluorescence microscope.

### Laser capture microdissection coupled mass spectrometry (LCM-MS)

Following the histology protocol above, 4µm serial sections were mounted onto MMI membrane slides (MMI, 50102) and stained with 0.04% Toluidine blue in 0.1M sodium acetate for 2 hours. Slides were washed in increasing serial ethanol washes and stored at 4°C. For laser dissection and collection, slides were loaded onto the MMI CellCut Laser Microdissection system (Molecular Machines & Industries). Using the MMI CellCut software and MMIs CapLift technology (MMI, 50204), regions of interest for the intervertebral disc (IVD) and bone were manually selected and collected as previously described^67,68^, using a laser focus of 400 μm, automated cutting at 2 mW laser power, at 40 μm/sec speed. >750,000 µm^2^ of tissue were collected per region, per *wt* and *lmx1ba^−/−^* fish at 9mpf. Captured specimens were stored at −80 °C and processed for liquid chromatography coupled tandem mass spectrometry.

### Liquid chromatography separation for mass spectrometry

Separation was performed on a Thermo Vanquish Neo UHPLC system configured with buffer A as 0.1% formic acid in water and buffer B as 0.1% formic acid in 80% acetonitrile. A specified injection volume was loaded on to the analytical column (IonOpticks Aurora series Rapid TS, 8cm x 75um ID, 1.7 um C18) kept at 35°C at an initial rate of 800 nl/min which was then dropped to 200 nl/min in 0.25. The separation was started during this time, with a gradient of 1% B to 5% B over this period. The next step was 5% B to 40% B over 20.25 minutes, 40% B to 100% B over 0.5 minutes before washing for 2 minutes at 100% B and dropping down to 1% B in 0.5 minute. The complete method time was 30 minutes. The analytical column was connected to a Thermo Orbitrap Astral mass spectrometer via a Thermo EasySpray Ion, with a nanospray voltage set at 1600 V and the ion transfer tube temperature set to 290°C.

### Mass spectrometry (MS) settings for DIA

Data was acquired in a data independent manner with an expected peak width of 6 seconds and a default charge state of 2. Full MS data was acquired in positive mode over a scan range of 350 to 1750 Th, with a resolution of 240,000, a custom normalised AGC target of 500 % and a maximum injection time of 10 ms for a single microscan. Fragmentation data was obtained according to a table of 200 m/z windows, with 3 Th m/z widths. The DIA window type was set to automatic, and window placement optimisation selected. These were acquired in the astral, with a normalised collision energy of 25%. The m/z range was 145-1450. A normalised AGC target of 500% was used with a maximum injection time of 7 ms for a single microscan. All data was collected in centroid mode with a loop control set to 0.6 s.

### Mass spectrometry analysis

Raw files were analysed using DIANN 2.3.2 using a “library-free” approach and default settings. Briefly, the SwissProt whole zebrafish proteome was downloaded and theoretical spectral library generated in “Prediction from FASTA” Mode with the following parameter: specific tryptic digestion, 1 missed cleavage, no variable modifications with N-terminal M excision and Carbamidomethyl (C) enabled as fixed modifications. Peptides were limited to 7-30aa in length with a precursor charge state between 1 and 4 and m/z range of 300 to 1800. Raw files were searched against this resultant spectral library using “Library search” mode with match between runs and “Gene” based proteotypicity with a 1% peptide level FDR and allowing auto-calibration of MS1 and MS2 accuracy, and scan window. Results were log2 transformed and median centred before analysing.

### Tissue preparation for GAG MSI

Sagittal histological sections of *wt* and *lmx1ba^−/−^* fish at 9mpf were prepared as above but unstained. CS disaccharide (CD002-5) and oligosaccharide (CS004), and HA (HA004) standards were acquired from Iduron (UK). Enzymes were applied to slides with a TM HTX3 (HTX Technologies LLC, NC, USA) as previously described^49^ and incubated in a pseudo-humidity chamber^69^. Chondroitinase ABC (9024-13-9; Merck, Germany) was applied for 2 hours. Matrix was applied and sample imaged. After acquisition, matrix was removed with 100% MeOH before sequential washing twice in 100% EtOH for two minutes, 96% EtOH for one minute, 70% EtOH for one minute and then 100% LC grade water (7732-18-5; Fisher Scientific) for two minutes twice. Heparinse (Hpase) I through III (P0735S, P0736S and P0737S, New England biolabs, MA, USA) was applied (1 U.ml^-1^ of each enzyme). After 14 hours, the sample was desiccated and keratanase II (KSase II (K0069, Tokyo Chemical Industry, Japan) 1 U.ml^-1^) applied for 2 hours before MSI comparing.

### Mass spectrometry for GAG MSI and data analysis

MSI analysis was undertaken using a tims-TOF fleX (Bruker Daltonics, Bremen, DE) as described in^49^. The MALDI matrix, 10 mg.ml-1 9-aminoacridine (92817; Sigma Aldrich, MO, USA), 0.1% formic acid (Z0797502 21; Sigma Aldrich) in 69.95% methanol and 29.95% LC grade water was applied using a TM HTX3 sprayer ^49^. Flex Imaging (Bruker Daltonics) software was used to set pixel size and imaging coordinates. Cohort images acquired at 20 μm pixel size and representative images at 5μm. MS/MS experiments were performed on standards or directly from tissue using CID (25-55 eV). Comparison of sample TIMS profiles to those of GAG standards was also used to aid identification.

SCILS lab (Bruker) environment was used to visualise datasets. All images were normalised to the total ion count (TIC). For TIMS profile analysis, Compass data analysis (Bruker) was used to select whole sections or regions of interest and mobillograms were subsequently generated for ions of interest in that region. Mobillograms were exported as .xy files and statistically analysed in R. Mobilities were normalised to maximum signal intensity (0-1). Quantification of isomers with TIMS was performed as described^50^. We describe ions based on disaccharide and sulphate composition as in^49^. For analysis of GAG composition and abundance in spines, signal was normalised to the TIC and mean pixel intensity measured across the whole spine for all GAG ions, which were summed into their GAG family and their values compared.

### 3D analysis of osteocyte lacunae

Segmentation and morphological analysis of osteocyte lacunae from SRCT images was performed as in^26^, albeit with an updated U-Net pixel classification model trained on example images taken from this study, and full segmentation and analysis workflow being conducted using the ModularImageAnalysis (MIA; version 1.7.22) plugin for Fiji^70^. This included an additional step incorporating application of the U-Net model using an implementation of the DeepImageJ plugin^71^. Weights for the new model were initialised using the final weights optimised as part of ^26^. For each fish, substacks of 400 slices over one IVD region were analysed.

### Swim behaviour analysis

Individual *wt* or *lmx1ba^−/−^* adults at 9 and 17mpf were placed in an 8 L tank (Techniplast) and recorded for 10 minutes from above at 15 frames per second. Fish were acclimated to the tank for 15 minutes before filming to reduce stress-induced behavioural artifacts. Tracking of fish location and body curvature per frame was performed using the MIA plugin for Fiji^70^.

Fish were segmented in each frame of the video using custom U-Net models, then tracked between frames with the TrackMate plugin for Fiji^72^. In each frame, a spline was fit along the length of the fish. From this spline curve, the local curvature of the fish was measured in correlation to the length of the background where right-sided bends had a maximum value of +1 and left-sided bends a maximum value of −1. For each video, 500 frames (∼33 seconds), including a minimum of 15 curvature events, were analysed.

### Statistics

Statistical analyses were performed using Graphpad Prism v.11.0.0. Error bars on all graphs represent the mean ± standard deviation. Sample sizes are based on a priori power calculations using previous data from adult zebrafish experiments performed in our lab.

## ACKNOWLEDGMENTS

The authors would like to acknowledge the Diamond Light Source for access and beamtime to the X-ray beamline I13-2 under proposal MG41910. We would also like to thank Dr Andre Rowe and Rabia Sevil for help with synchrotron scanning at DLS. We would like to thank Dr Qiao Tong for her help with μCT scanning. The proteomics was performed at the Biological Mass Spectrometry Facility at University of Manchester, with the assistance of Stacey Warwood. We thank the staff of the Wolfson Bioimaging Facility for imaging support, especially Dr Dani Franchini for her help with developing the swim analysis MIA platform. We also thank Mathew Green and technical staff from the University of Bristol’s Animal Scientific Unit (ASU) for providing zebrafish husbandry. We further thank Debbie Martin from the University of Bristol histology facility for help with histological processing.

## AUTHOR CONTRIBUTIONS

J.J.M. and C.L.H. conceptualised the study and designed experiments. J.J.M. performed most of the experiments. J.C. performed the mass spectrometry from laser dissection, A.D. performed the GAG analysis and mass spectrometry, F.B. performed µCT scanning, E.N, F.B. J.J.M. E.R.J. and C.L.H. performed SRCT scanning, S.C. developed the lacunae analysis and swim analysis workflows. J.J.M. performed statistical analysis and made all figures. J.J.M. and C.L.H wrote the first draft of the manuscript, and all authors made intellectual contributions and assisted in manuscript editing.

## DATA AVAILABILITY

All raw data will be available via a DOI at data.bris.ac.uk upon article acceptance.

## REFERENCES

1 Curtiss, J. & Heilig, J. S. DeLIMiting development. Bioessays 20, 58–69 (1998).

2 Hobert, O. & Westphal, H. Functions of LIM-homeobox genes. Trends in genetics 16, 75–83 (2000).

3 Suleiman, H. et al. The podocyte-specific inactivation of Lmx1b, Ldb1 and E2a yields new insight into a transcriptional network in podocytes. Developmental biology 304, 701–712 (2007).

4 Feenstra, J. M. et al. Detection of genes regulated by Lmx1b during limb dorsalization. Development, growth & differentiation 54, 451–462 (2012).

5 Gu, W. X. & Kania, A. Identification of genes controlled by LMX1B in E13. 5 mouse limbs. Developmental Dynamics 239, 2246–2255 (2010).

6 Krawchuk, D. & Kania, A. Identification of genes controlled by LMX1B in the developing mouse limb bud. Developmental Dynamics 237, 1183–1192 (2008).

7 Haro, E. et al. Lmx1b-targeted cis-regulatory modules involved in limb dorsalization. Development 144, 2009–2020 (2017).

8 Bian, Q., Wang, Y.-J., Liu, S. F. & Li, Y.-P. Osteoarthritis: genetic factors, animal models, mechanisms, and therapies. Front Biosci (Elite Ed) 4, 74–100 (2012).

9 Zhang, X. et al. Global, regional, and country-specific lifetime risks of osteoarthritis, 1990–2021: a systematic analysis for the global burden of disease study 2021. Global Health Research and Policy 10, 29 (2025).

10 Neogi, T. The epidemiology and impact of pain in osteoarthritis. Osteoarthritis and cartilage 21, 1145–1153 (2013).

11 Nicolae, C. et al. Abnormal collagen fibrils in cartilage of matrilin-1/matrilin-3-deficient mice. Journal of biological chemistry 282, 22163–22175 (2007).

12 Saklatvala, J. Does decorin stabilize the extracellular matrix of articular cartilage and slow the progression of osteoarthritis? Osteoarthritis and Cartilage (2021).

13 Li, P. et al. Mice lacking the matrilin family of extracellular matrix proteins develop mild skeletal abnormalities and are susceptible to age-associated osteoarthritis. International journal of molecular sciences 21, 666 (2020).

14 Miyamoto, Y. et al. A functional polymorphism in the 5′ UTR of GDF5 is associated with susceptibility to osteoarthritis. Nature genetics 39, 529–533 (2007).

15 Barreto, G. et al. Lumican is upregulated in osteoarthritis and contributes to TLR4-induced pro-inflammatory activation of cartilage degradation and macrophage polarization. Osteoarthritis and Cartilage 28, 92–101 (2020).

16 Melrose, J. et al. Fragmentation of decorin, biglycan, lumican and keratocan is elevated in degenerate human meniscus, knee and hip articular cartilages compared with age-matched macroscopically normal and control tissues. Arthritis research & therapy 10, 1–10 (2008).

17 Kim, K. et al. Transcription factor Lmx1b negatively regulates osteoblast differentiation and bone formation. International Journal of Molecular Sciences 23, 5225 (2022).

18 Wang, C. et al. Single-cell RNA sequencing analysis of human embryos from the late Carnegie to fetal development. Cell & Bioscience 14, 118 (2024).

19 Morris, J. A. et al. An atlas of genetic influences on osteoporosis in humans and mice. Nature genetics 51, 258–266 (2019).

20 Tachmazidou, I. et al., doi:10.1101/453530 (2018).

21 Chen, C., Kong, D., Wang, P., Li, M. & Gui, R. Genetic polymorphisms of LMX1B and MLXIP are associated with hip osteoarthritis in the Chinese population. Biomarkers in Medicine 18, 695–702 (2024).

22 Boer, C. G. et al. Deciphering osteoarthritis genetics across 826,690 individuals from 9 populations. Cell 184, 4784–4818. e4717 (2021).

23 Sweeney, E., Fryer, A., Mountford, R., Green, A. & McIntosh, I. Nail patella syndrome: a review of the phenotype aided by developmental biology. Journal of medical genetics 40, 153–162 (2003).

24 Tachmazidou, I. et al. Identification of new therapeutic targets for osteoarthritis through genome-wide analyses of UK Biobank data. Nature genetics 51, 230–236 (2019).

25 Moss, J. J., Neal, C. R., Kague, E., Lane, J. D. & Hammond, C. L. Characterisation of lmx1b paralogues in zebrafish reveals divergent roles in skeletal, kidney and muscle development. Biology Open 14, bio062038 (2025).

26 Kague, E. et al. 3D assessment of intervertebral disc degeneration in zebrafish identifies changes in bone density that prime disc disease. Bone research 9, 39 (2021).

27 Park, J.-S. et al. Characterization of zebrafish nucleus pulposus in development and aging: A fish model for probing human intervertebral disc degeneration and regeneration. bioRxiv, 2025.2010. 2020.682618 (2025).

28 Kague, E., Larraz-Prieto, B., Raele, R., Moss, J. & Hammond, C. Targeted modulation of phosphate and lipid metabolism reduces ligament mineralization in col9a1b knockout models. (2025).

29 Spoorendonk, K. M., et al. Retinoic acid and Cyp26b1 are critical regulators of osteogenesis in the axial skeleton. (2008).

30 Pogoda, H.-M. et al. Direct activation of chordoblasts by retinoic acid is required for segmented centra mineralization during zebrafish spine development. Development 145, dev159418 (2018).

31 Martini, A. et al. Deformity or variation? Phenotypic diversity in the zebrafish vertebral column. Journal of Anatomy 243, 960–981 (2023).

32 Kim, K., Han, J. E., Lee, K.-B. & Kim, N. LIM homeobox transcription factor 1-β expression is upregulated in patients with osteolysis after total ankle arthroplasty and inhibits receptor activator of nuclear factor-κB ligand-induced osteoclast differentiation in vitro. Journal of Bone Metabolism 29, 165 (2022).

33 López-Cuevas, P., Deane, L., Yang, Y., Hammond, C. L. & Kague, E. Transformed notochordal cells trigger chronic wounds in zebrafish, destabilizing the vertebral column and bone homeostasis. Disease Models & Mechanisms 14, dmm047001 (2021).

34 Ricard-Blum, S., Baffet, G. & Théret, N. Molecular and tissue alterations of collagens in fibrosis. Matrix Biology 68, 122–149 (2018).

35 Swann, D., Silver, F., Slayter, H., Stafford, W. & Shore, E. The molecular structure and lubricating activity of lubricin isolated from bovine and human synovial fluids. Biochemical Journal 225, 195–201 (1985).

36 Han, S. Osteoarthritis year in review 2022: biology. Osteoarthritis and cartilage 30, 1575–1582 (2022).

37 Karamanos, N. K. et al. Proteoglycan chemical diversity drives multifunctional cell regulation and therapeutics. Chemical reviews 118, 9152–9232 (2018).

38 Chen, J. et al. Proteoglycans and glycosaminoglycans in stem cell homeostasis and bone tissue regeneration. Frontiers in Cell and Developmental Biology 9, 760532 (2021).

39 Hascall, V. C. & Sajdera, S. W. Physical properties and polydispersity of proteoglycan from bovine nasal cartilage. Journal of Biological chemistry 245, 4920–4930 (1970).

40 Ren, K., Zhao, F., Tzaneti, N. K., Kaper, H. J. & Sharma, P. K. Glycosaminoglycan depletion lowers the crack resistance of articular cartilage under impact loading. Journal of the mechanical behavior of biomedical materials 170, 107122 (2025).

41 Hua, R. et al. Biglycan and chondroitin sulfate play pivotal roles in bone toughness via retaining bound water in bone mineral matrix. Matrix Biology 94, 95–109 (2020).

42 Soares da Costa, D., Reis, R. L. & Pashkuleva, I. Sulfation of glycosaminoglycans and its implications in human health and disorders. Annual review of biomedical engineering 19, 1–26 (2017).

43 Lauder, R. M., Huckerby, T. N., Brown, G. M., Bayliss, M. T. & Nieduszynski, I. A. Age-related changes in the sulphation of the chondroitin sulphate linkage region from human articular cartilage aggrecan. Biochemical Journal 358, 523–528 (2001).

44 Fisher, L. W. et al. Proteoglycans of developing bone. Journal of Biological Chemistry 258, 6588–6594 (1983).

45 Plaas, A. H., West, L. A., Wong-Palms, S. & Nelson, F. R. Glycosaminoglycan sulfation in human osteoarthritis: disease-related alterations at the non-reducing termini of chondroitin and dermatan sulfate. Journal of Biological Chemistry 273, 12642–12649 (1998).

46 Lark, M. W., Bayne, E. K. & Lohmander, L. S. Aggrecan degradation in osteoarthritis and rheumatoid arthritis. Acta Orthopaedica Scandinavica 66, 92–97 (1995).

47 Reynard, L. N., Ratnayake, M., Santibanez-Koref, M. & Loughlin, J. Functional characterization of the osteoarthritis susceptibility mapping to CHST11—a bioinformatics and molecular study. PLoS One 11, e0159024 (2016).

48 Watson, H. J. et al. Genome-wide association study identifies eight risk loci and implicates metabo-psychiatric origins for anorexia nervosa. Nature genetics 51, 1207–1214 (2019).

49 Devlin, A., Green, F. & Takats, Z. Mass spectrometry imaging with trapped ion mobility spectrometry enables spatially resolved chondroitin, dermatan, and hyaluronan glycosaminoglycan oligosaccharide analysis in situ. Analytical Chemistry 96, 17969–17977 (2024).

50 Anthony Devlin, F. G. Towards spatially resolved disaccharide analysis: localisation of different populations of chondroitin sulphate disaccharide isomers liberated directly from tissue sections using trapped ion mobility spectrometry mass spectrometry (TIMS-MS). BioRxiv (2026).

51 Hayes, A. J. et al. Spinal deformity in aged zebrafish is accompanied by degenerative changes to their vertebrae that resemble osteoarthritis. PloS one 8, e75787 (2013).

52 Boskey, A. L. et al. Examining the relationships between bone tissue composition, compositional heterogeneity, and fragility fracture: a matched case-controlled FTIRI study. Journal of Bone and Mineral Research 31, 1070–1081 (2016).

53 McGowan, L. M. et al. Wnt16 elicits a protective effect against fractures and supports bone repair in zebrafish. Journal of Bone and Mineral Research Plus 5, e10461 (2021).

54 Tiede-Lewis, L. M. & Dallas, S. L. Changes in the osteocyte lacunocanalicular network with aging. Bone 122, 101–113 (2019).

55 Carpentier, V. T. et al. Increased proportion of hypermineralized osteocyte lacunae in osteoporotic and osteoarthritic human trabecular bone: implications for bone remodeling. Bone 50, 688–694 (2012).

56 Fiedler, I. A. et al. Severely impaired bone material quality in chihuahua zebrafish resembles classical dominant human osteogenesis imperfecta. Journal of Bone and Mineral Research 33, 1489–1499 (2018).

57 Goldring, S. R. & Goldring, M. B. Changes in the osteochondral unit during osteoarthritis: structure, function and cartilage–bone crosstalk. Nature Reviews Rheumatology 12, 632–644 (2016).

58 Loeser, R. F., Goldring, S. R., Scanzello, C. R. & Goldring, M. B. Osteoarthritis: a disease of the joint as an organ. Arthritis and rheumatism 64, 1697 (2012).

59 Yang, Y. et al. Tuning collective behaviour in zebrafish with genetic modification. PLoS Computational Biology 20, e1012034 (2024).

60 Baird, D. A. et al. Investigation of the relationship between susceptibility loci for hip osteoarthritis and dual x-ray absorptiometry–derived hip shape in a population-based cohort of perimenopausal women. Arthritis & Rheumatology 70, 1984–1993 (2018).

61 Wilkinson, J. M. & Zeggini, E. The genetic epidemiology of joint shape and the development of osteoarthritis. Calcified tissue international 109, 257–276 (2021).

62 Aleström, P. et al. Zebrafish: Housing and husbandry recommendations. Laboratory Animals 54, 213–224 (2020).

63 Knopf, F. et al. Bone regenerates via dedifferentiation of osteoblasts in the zebrafish fin. Developmental cell 20, 713–724 (2011).

64 van den Berg, M. C., et al. Proteolytic and opportunistic breaching of the basement membrane zone by immune cells during tumor initiation. Cell reports 27, 2837–2846. e2834 (2019).

65 Garcia, J. et al. Sheath cell invasion and trans-differentiation repair mechanical damage caused by loss of caveolae in the zebrafish notochord. Current Biology 27, 1982–1989. e1983 (2017).

66 Dane, E. T. & Herman, D. L. Haematoxylin-phloxine-Alcian blue-orange G differential staining of prekeratin, keratin and mucin. Stain Technology 38, 97–101 (1963).

67 Herrera, J. A. et al. Laser capture microdissection coupled mass spectrometry (LCM-MS) for spatially resolved analysis of formalin-fixed and stained human lung tissues. Clinical Proteomics 17, 24 (2020).

68 Revell, C. K. et al. Modeling collagen fibril self-assembly from extracellular medium in embryonic tendon. Biophysical Journal 122, 3219–3237 (2023).

69 Angel, P. M., Norris-Caneda, K. & Drake, R. R. In situ imaging of tryptic peptides by MALDI imaging mass spectrometry using fresh-frozen or formalin-fixed, paraffin-embedded tissue. Current protocols in protein science 94, e65 (2018).

70 Cross, S. J., Fisher, J. D. & Jepson, M. A. ModularImageAnalysis (MIA): Assembly of modularised image and object analysis workflows in ImageJ. Journal of microscopy 296, 173–183 (2024).

71 Gómez-de-Mariscal, E. et al. DeepImageJ: A user-friendly environment to run deep learning models in ImageJ. Nature methods 18, 1192–1195 (2021).

72 Tinevez, J.-Y. et al. TrackMate: An open and extensible platform for single-particle tracking. Methods 115, 80–90 (2017).

