## Supplemental Figures for "Loss of *lmx1ba* drives premature osteoarthritis through disruption of skeletal homeostasis"

### Supporting information listing

| Item | Type | Title |
|---|---|---|
| Supplemental Figure 1 | Figure | Spinal morphology is altered in adult *lmx1ba* mutants. |
| Supplemental Figure 2 | Figure | Severity of spinal deformities in *lmx1ba* mutants is comparable to much older *wt* fish. |
| Supplemental Figure 3 | Figure | Vertebral bone mineralisation is delayed in *lmx1ba*$^{-/-}$ larvae but recovers by 14dpf whilst vertebral segmentation remains unaffected. |
| Supplemental Figure 4 | Figure | Minor morphological alterations in spine shape in *lmx1ba* mutants are maintained at 3mpf. |
| Supplemental Figure 5 | Figure | Heparan sulphate (HS) and keratan sulphate (KS) distribution is unaltered in adult *lmx1ba*$^{-/-}$ spines. |
| Supplemental Figure 6 | Figure | Examples of sample acquisition steps from laser dissection for mass spectrometry analysis of different tissue regions. |

### Supplemental Figures and Figure Legends

#### Supplemental Figure 1

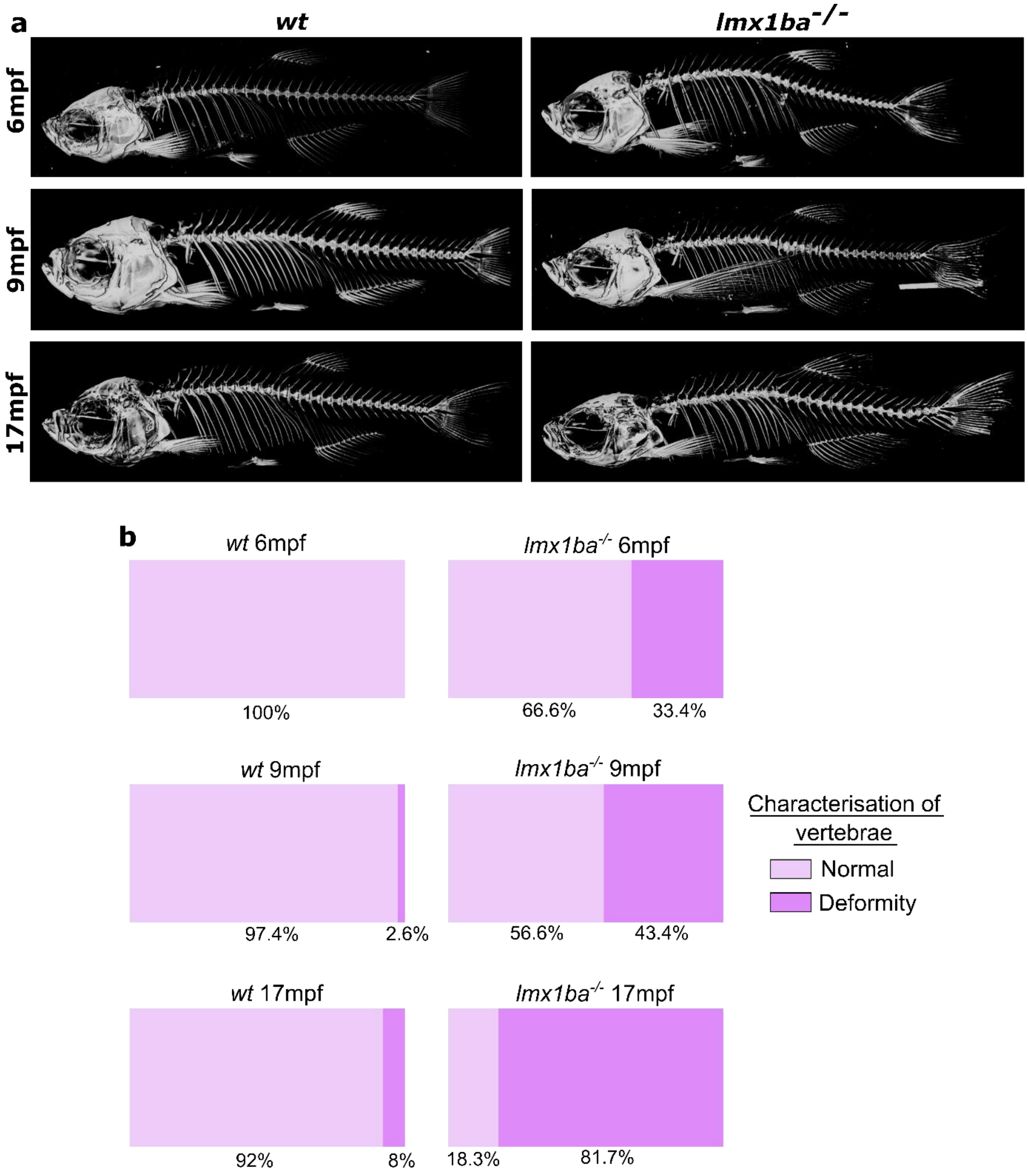

**Supplemental Figure 1 – Spinal morphology is altered in adult *lmx1ba* mutants.**

**(A)** 3D rendering from µCT images of *wt* and *lmx1ba*$^{-/-}$ fish at 6, 9 and 17mpf. **(B)** Stacked bar charts showing percentage of different vertebral deformities by age and genotype. Calculated as the average percentage of vertebrae with or without a deformity from the pre-caudal spine to the caudal fin spine per age. N = 3 at 6mpf; N = 5 at 9 and 17mpf.

**Supplemental Figure 2**

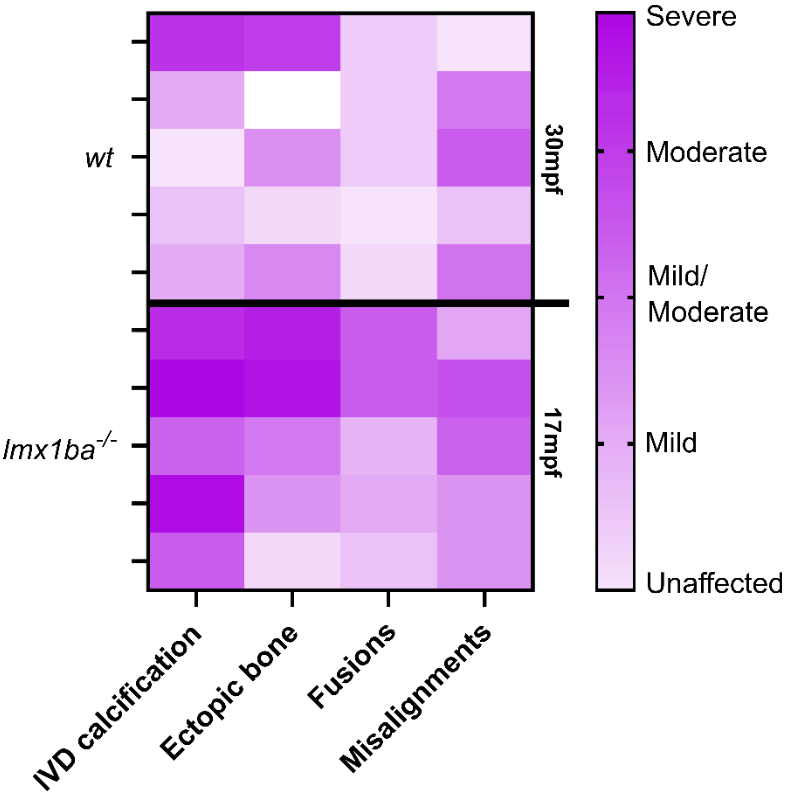


**Supplemental Figure 2 – Severity of spinal deformities in *lmx1ba* mutants is comparable to much older *wt* fish.**

Heat map showing the degree of severity of different spinal deformities between *wt* fish at 30mpf and *lmx1ba*$^{-/-}$ fish at 17mpf. N = 5 per group.

### Supplemental Figure 3

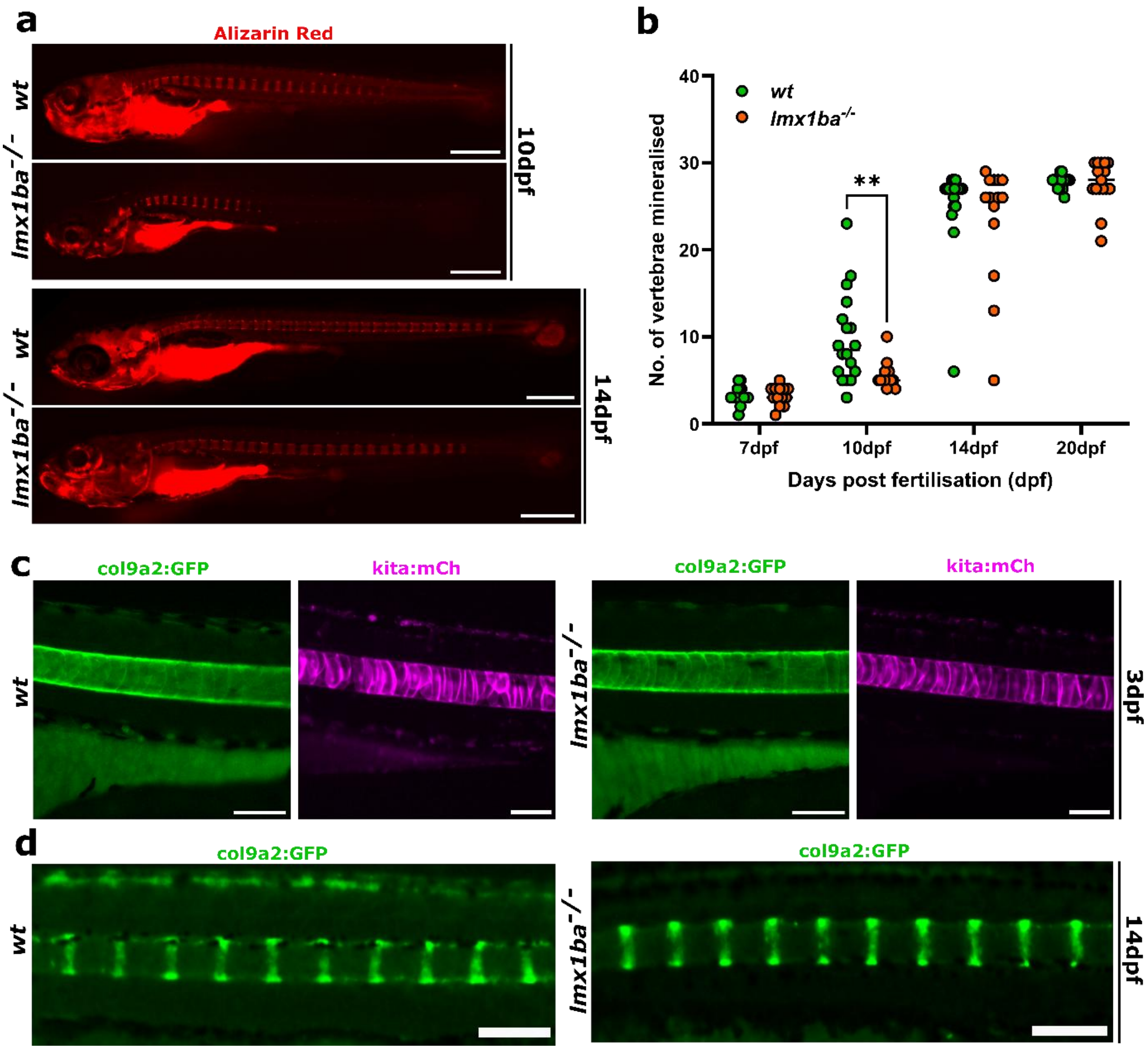

**Supplemental Figure 3 – Vertebral bone mineralisation is delayed in *lmx1ba*$^{-/-}$ larvae but recovers by 14dpf whilst vertebral segmentation remains unaffected.**

**(A)** Representative lateral stereomicroscope images of *wt* and *lmx1ba*$^{-/-}$ larvae at 10dpf and 14dpf following live staining with Alizarin red for 1 hour. Scale bars = 500µm for all. **(B)** Graph showing number of mineralized vertebrae per fish at each time point. N = 18 for *wt* and 14-17 for *lmx1ba*$^{-/-}$. A mixed-effects ANOVA test was performed where ***P*=.0087. **(C)** Notochord images of *col9a2:GFP* (green) and *kita:mCherry* (magenta) *wt* and *lmx1ba*$^{-/-}$ larvae at 3dpf. Scale bars = 100µm. **(D)** Spinal images of *col9a2:GFP* (green) *wt* and *lmx1ba*$^{-/-}$ larvae at 14dpf. Scale bars = 200µm.

**Supplemental Figure 4**

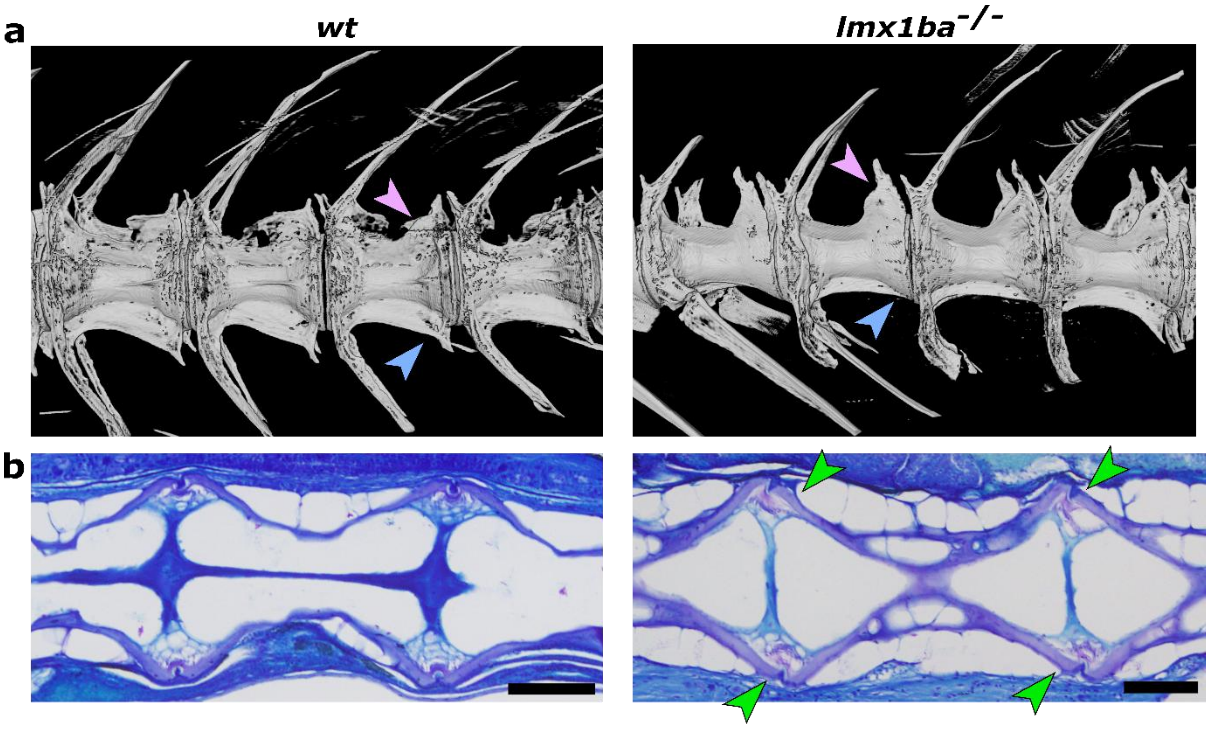


**Supplemental Figure 4 – Minor morphological alterations in spine shape in *lmx1ba* mutants are maintained at 3mpf.**

**(A)** 3D volumetric rendering of synchrotron radiation-based µCT images of *wt* and *lmx1ba*$^{-/-}$ spines at 3mpf. Arrowheads show morphological differences in npstz (pink) and hpstz (blue) formation. N = 3 per group. **(B)** Histological sections of zebrafish IVDs at 3mpf stained with Toluidine blue. Green arrowheads show regions of vertebral misalignments. Scale bars = 50 µm. N = 5 per group. Npstz = neural postzygapophyses; hpstz = hemal postzygapophyses.

### Supplemental Figure 5

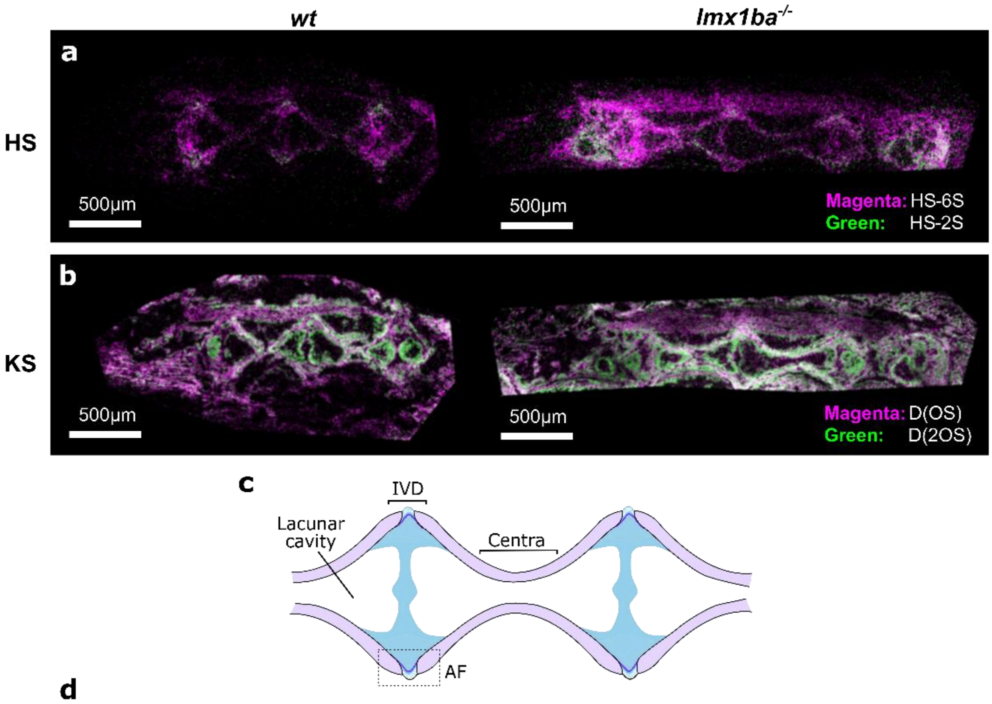


**d**

| Region | | HS | | KS | |
|---|---|---|---|---|---|
| | | **HS-2S / %** | **P-value** | **KS D(OS) / %** | **P-value** |
| Whole section | *wt* | 2.4 +/- 1.4 | 0.14 | 61.2 +/- 2.8 | 0.53 |
| | *lmx1ba$^{-/-}$* | 5.4 +/- 1.7 | | 62.8 +/- 2.3 | |
| Whole Spine | *wt* | 10.5 +/- 3.3 | 0.66 | 67.2 +/- 0.8 | 0.64 |
| | *lmx1ba$^{-/-}$* | 12.1 +/- 3.6 | | 64.4 +/- 7.1 | |
| Centra | *wt* | 7.9 +/- 1.2 | 0.26 | 54.1 +/- 0.1 | 0.003 |
| | *lmx1ba$^{-/-}$* | 9.2 +/- 0.3 | | 49.9 +/- 0.3 | |
| AF | *wt* | 10.4 +/- 0.8 | 0.22 | 54.7 +/- 4.4 | 0.19 |
| | *lmx1ba$^{-/-}$* | 11.4 +/- 0.3 | | 60.4 +/- 0.8 | |
| IVD | *wt* | n/a | n/a | 50.2 +/- 4.6 | 0.24 |
| | *lmx1ba$^{-/-}$* | n/a | | 56.2 +/- 4.1 | |

**Supplemental Figure 5 – Heparan sulphate (HS) and keratan sulphate (KS) distribution is unaltered in adult *lmx1ba*$^{-/-}$ spines.**

Trapped ion mobility spectrometry mass spectrometry imaging (TIMS-MSI) of **(A)** HS DP2s and **(B)** KS DP2s for *wt* and *lmx1ba*$^{-/-}$ fish. N = 2 per group. **(C)** Schematic showing location of vertebral regions measured in **(D)** Table showing relative abundance of detected HS and KS disaccharides. Students T-test performed between *wt* and *lmx1ba*$^{-/-}$ fish of the same region. HS = heparan sulphate; KS = keratan sulphate. n/a values appear where HS signal could not be detected.

**Supplemental Figure 6**

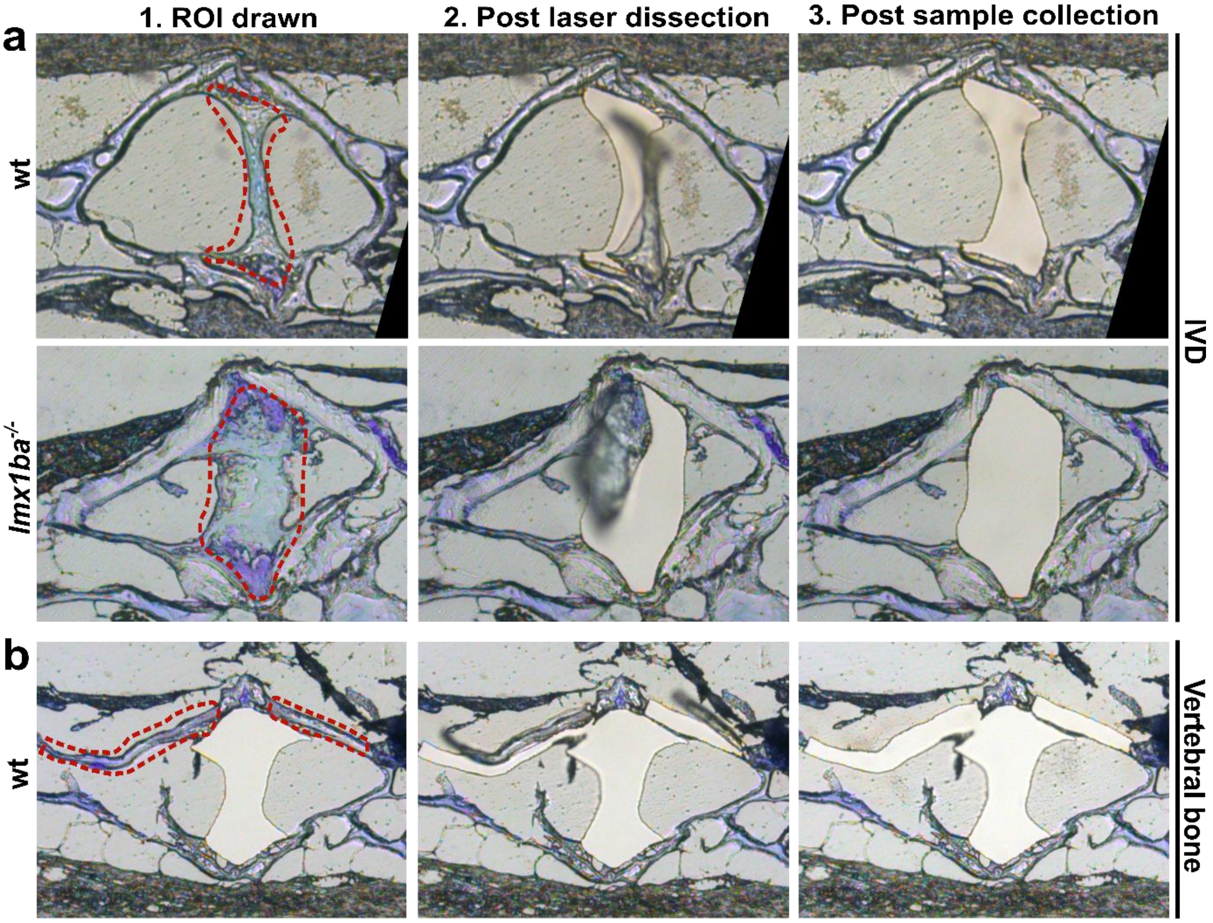


**Supplemental Figure 6 – Examples of sample acquisition steps from laser dissection for mass spectrometry analysis of different tissue regions.**

Confocal images of toluidine blue stained vertebrae from 9mpf *wt* and *lmx1ba*$^{-/-}$ histological sections. Red dotted line shows examples of **(A)** IVD or **(B)** vertebral bone regions of interest (ROI) laser dissected by the MMI CellCut Laser Microdissection system and sequentially collected for mass spectrometry analysis.
